# Predicting cell-state-specific gene regulation from DNA across species

**DOI:** 10.64898/2026.10.19.760026

**Authors:** Hanqing Liu, Sheng Mao, Tristan Gallent, Yishuang Chen, Zhiwen Yan, Ian Vogel, Lisa Traunmüller, Ruochi Zhang, Jason D Buenrostro, Jingtian Zhou, Michael E Greenberg

## Abstract

DNA encodes the cell’s phenotype, but the same DNA sequence is interpreted differently in every cell type, developmental stage, and activity state. Existing deep-learning models solve only half of this puzzle: sequence-to-function models read DNA but treat cell state as a fixed label, while single-cell foundation models represent cell state but not sequence. We developed Bolero, a sequence-to-function model whose parameters condition on atlas-scale cell-state embeddings and additional biological variables to predict cell-state-specific chromatin accessibility and transcript abundance from DNA. Trained on 10.7 million cells from 33 datasets across six mammals, Bolero matched or exceeded specialised models and resolved regulatory differences between closely related cell states, including cortical regional and developmental gradients. Extrapolating cell-state embeddings extended Bolero’s predictions to cell states that are absent from training. Cell-state conditioning facilitated tracing of the effects of non-coding variants to the cell types in which they act. Using an AP-1 activity score as a further continuous conditioning axis, Bolero predicted activity-regulated chromatin changes without requiring actual stimulation data. Across human cell types, we reconstructed the cooperative grammar of motif identity and spacing underlying predicted AP-1 responses. Bolero further predicted cell-type-specific chromatin accessibility and AP-1 responses in species excluded from training, extending comparative genomics from sequence conservation to predicted functional conservation. Across 84 mammals, we found that predicted AP-1 responses were less conserved than chromatin accessibility at cis-regulatory elements, driven by the human-specific sequence variants on AP-1 and cooperating motifs. By combining reference genomes with reference cell atlases, Bolero allowed genome sequences, either native or mutated, to be queried across cell states, perturbations, and different species for their chromatin and expression signatures.

## Introduction

Organismal function is mediated by hundreds of cell types and a continuous spectrum of developmental, physiological, and stimulus-induced states, each engaging its own cis-regulatory element-driven gene expression program. Multimodal single-cell gene expression reference atlases have catalogued this diversity across whole organs^1,2^, developmental trajectories^3,4^, and dozens of mammals^5^. However, it is impractical and costly for experimental atlases to resolve the full combinatorial space spanned by cell type, stimulus, developmental stage, and genetic background. The most consequential downstream applications of these atlases, including non-coding variant effect interpretation, perturbation prediction, and cross-species comparison, require predictions for states that these atlases have not captured. Single-cell foundation models^6–11^ can help address part of this gap by learning extrapolable embeddings of cell state. Yet they operate on a fixed gene-expression feature space and cannot directly interpret regulatory information encoded in DNA or predict mutation effects on gene expression and cell states. Available sequence-to-function models^12–18^ offer complementary capabilities, including flexible DNA input and base-pair-resolution attribution. But they lack an explicit representation of cell state and do not generalise to cell states absent from the training data.

We developed Bolero, a model that learns how sequence is read as a function of the cell state’s position in an embedding space and predicts chromatin accessibility and transcript abundance from DNA at any point in that space. This is achieved by conditioning the sequence-to-function model Borzoi^17^ on an atlas-scale cell-state embedding plus orthogonal conditioning variables, such as developmental age and the effect of external stimuli. Cell state is therefore an input the user sets, not one of a fixed set of categories chosen at training time (Fig. 1a). Trained on Bolero-10M, a 10.7-million-cell single-cell epigenome and multiomics atlas spanning mammalian brain and body, development and ageing, and evolution, Bolero achieved state-of-the-art accuracy on cell-state-specific regulation and sequence variant-effect prediction and generalised to cell states unseen during training. With an AP-1 activity score as a further conditioning axis, Bolero predicted which regulatory elements are activated or repressed in behaviour-paired paradigms held out from training. Attribution analysis and in silico motif insertion then identified the cooperating transcription factors and motif spacings that shape the cell-type specificity of the AP-1 response. Applied to mammalian genomes, Bolero predicted AP-1-responsive candidate cis-regulatory elements (cCREs), including human-specific candidates, without requiring stimulus-evoked data from humans. Comparing these predictions across species showed that inducible AP-1 responses in basal-ganglia neurons are less conserved than the chromatin accessibility of the same elements. Changes in AP-1 and cooperating transcription-factor motifs partly explained these predicted differences in responsiveness. Bolero provides a general and extensible way to identify cell-state-specific regulatory codes by converting a reference atlas from a cell catalogue into a model of how the genome gives rise to cell-state-specific gene regulation.

**Figure 1.**
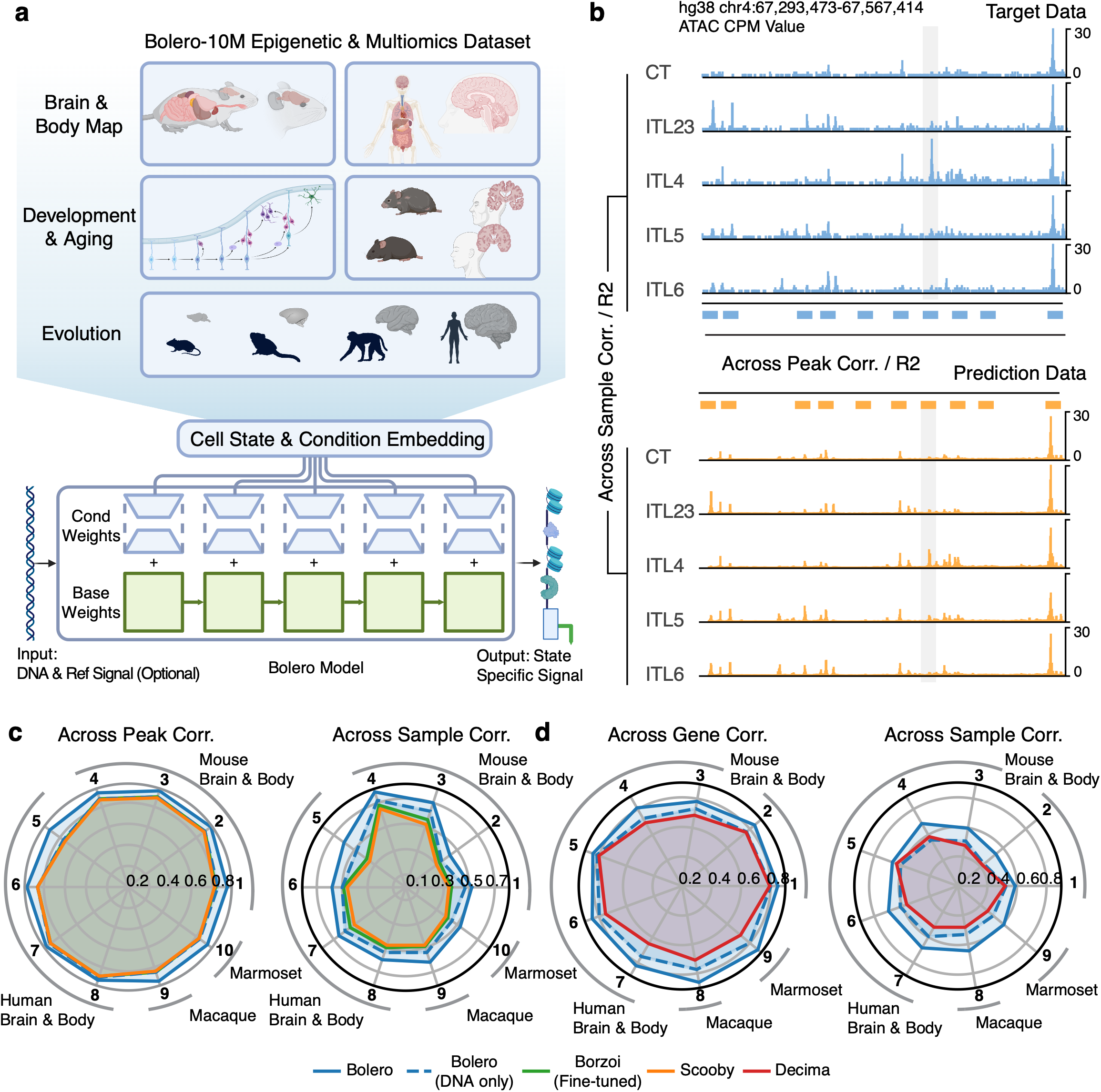
Bolero is a cell-state-conditioned sequence-to-function model. **a,** Overview schematic: the Bolero-10M atlas spans brain and body, development and ageing, and evolution (top); one-hot DNA (524,288 bp) and a per-cell-state embedding condition the Borzoi model to predict cell-state-resolved chromatin accessibility (ATAC, bottom). **b,** Measured (blue) and predicted (orange) ATAC tracks at an example locus across five cortical excitatory subclasses, with called peaks marked between the two track blocks; bracket labels denote the across-sample (down the subclasses) and across-peak evaluation axes. **c,d,** Radar plots of prediction accuracy across features (left; peaks in c, genes in d) or across samples (right) for chromatin accessibility (c) and for gene expression (d). Each spoke is a benchmark dataset and the radius is the Pearson correlation, coloured by model (DNA-only Bolero dashed, representing Bolero without bulk signal as input). The full benchmark dataset names are in Extended Data Fig. 1.

### A cell-state-conditioned sequence-to-function model trained on a 10M-cell atlas

Bolero is a sequence-to-function model with two inputs: a DNA sequence and a cell-state embedding (a compact numerical summary of a cell state, computed from that cell state’s single-cell measurements). An optional set of further conditioning variables, which describe the sample metadata beyond its cell-state embedding, can be supplied alongside them. Bolero outputs the chromatin accessibility and transcript abundance expected for that cell state (Fig. 1a, Extended Data Fig. 1a). The DNA is read by the Flashzoi variant of the Borzoi model^17,19^, whose weights are fixed and shared by every cell state. Bolero uses the cell-state embedding to generate a small set of extra weights and adds them to the fixed weights at every layer. Each cell state therefore gets its own effective network, while what Borzoi has learned about DNA stays common to all of them. The conditioning acts only through these added weights and does not depend on the architecture of the backbone, so the same construction can be applied to other sequence-to-function models, such as ChromBPNet^16^ or AlphaGenome^18^, to give them cell-state resolution. Because the cell state changes only the weights, every prediction can be attributed back completely to DNA sequence rather than cell embedding, enabling model interpretation in the same way as a DNA-only model like Borzoi. Developmental age, transcription factor activity, and species enter the same way, as further inputs trained jointly with the embedding, so a single model can be asked how one sequence behaves at a different age, under a different stimulus, or in another mammalian species.

To learn cell-state-resolved gene regulation at scale, we assembled Bolero-10M, a unified single-cell epigenome and multiomics atlas of 10.7 million cells from 33 datasets across six mammals (Fig. 1a; Supplementary Table 1). Bolero-10M spans cell types across brain and body, development and ageing, and evolutionary divergence. Single-cell embeddings were generated based on integrated chromatin and expression profiles. Conditioning the model with the continuous embedding space allows training to gain information from neighboring cell states. To ensure enough signal-to-noise, we merged single cells with similar embeddings into pseudobulk profiles with uniform sequencing depth, used as the unit samples for model training.

Bolero predictions reproduced both the distribution of accessibility along the DNA sequence and differences between cell states (Fig. 1b). To benchmark performance systematically, we compared Bolero against fine-tuned Borzoi and Scooby^20^ on 10 representative datasets spanning four species, evaluated on held-out genomic regions using the train/test split of Borzoi (Fig. 1c, d, Extended Data Fig. 1b,c). Bolero was run in its signal-aware mode, which supplies one reference chromatin accessibility profile that is identical for every cell state. It carries information about the scale of the predicted values and no information about which state is being predicted (see Methods). On per-sample, across-peak correlation, the axis on which existing sequence-to-function models already perform well, Bolero matched the strongest baseline (Borzoi), confirming that the new architecture preserves genomic resolution (Fig. 1c). On across-sample correlation, which measures whether cell states are distinguishable from each other, Bolero substantially outperformed every baseline in every benchmark dataset (Fig. 1c). Cell-state resolution gains therefore come from the embedding, and the conditional architecture is what makes it possible. Withholding that reference ATAC signal and predicting from DNA alone left both results standing across all 10 datasets: across-peak correlation was at or marginally above Borzoi’s, and across-sample correlation still exceeded every baseline (dashed, Fig. 1c; Supplementary Table 2). The same architecture also predicted cell-state-resolved gene expression, where Bolero likewise outperformed Decima^21^ on across-sample correlation (Fig. 1d, Extended Data Fig. 1c; Supplementary Table 3). The advantage was largest at the most cell-state-specific peaks and genes (Extended Data Fig. 2a-d). Bolero models trained jointly on all human or all mouse datasets (WholeHuman and WholeMouse) matched the per-dataset accuracy of Bolero models trained separately on each dataset (Extended Data Fig. 2e), allowing us to avoid per-dataset specialisation.

To test how far these gains depend on our design choices, we asked how accuracy scales with training data and with model size, and whether every part of the architecture is needed. Across-sample correlation rose as the model saw more training examples, each one a genomic region paired with a cell state, and then plateaued, showing that training had converged for the data budget and model sizes tested (Extended Data Fig. 3a). Model size behaved the same way. Accuracy improved as trainable parameters were added, up to roughly 1 × 10⁹, and then plateaued, with the largest models we tested giving no consistent further benefit (Extended Data Fig. 3b). This suggests that for the architectures and sizes we tested, Bolero is already making full use of the training data. Conditioning every layer of the network outperformed conditioning a single layer (Extended Data Fig. 3c), so the cell state has to shape the network at every depth, not only at its output. Five different ways of computing the cell-state embedding performed comparably (Extended Data Fig. 3d), which suggests the resolution gain mostly comes from having a cell-state representation rather than the method used to build it.

### Bolero predicts spatial and temporal regulation, including in held-out cell states

Cell-type identity, on which existing sequence-to-function models are usually evaluated, is a coarse axis of regulatory variation. Within a single cortical cell type or lineage, chromatin accessibility varies systematically along spatial axes (cortical region) and temporal axes (developmental stage), and these within-cell-type differences carry the biological signal relevant to disease, organismal development, and evolution. We next tested whether a sequence model can resolve this finer structure and extrapolate to cell states it has never seen.

We first asked whether Bolero captures the spatial regulatory variation within a single cortical cell type. In the mouse cortex, L2/3 intratelencephalic (IT) excitatory neurons differ in chromatin accessibility across brain regions^22–24^. Bolero’s predictions reproduced the regional chromatin accessibility pattern of L2/3 IT neurons across 21 brain regions (Fig. 2a). Predicted regional gradients also matched the measured ones at three representative loci, including *Cspp1*, *Fscn1*, and *Kcnh7*, mapped onto the mouse brain common coordinate framework^25^ (Fig. 2a, b). Across cell-subclass differentially accessible regions (DARs) and brain-region DARs, Bolero’s median across-sample correlation was 0.59 and 0.62, compared with 0.38 and 0.41 for Scooby, an improvement of about 50% (Fig. 2c). Bolero therefore resolves not just discrete cell types but also the within-type spatial gradients along which they are organised.

**Figure 2.**
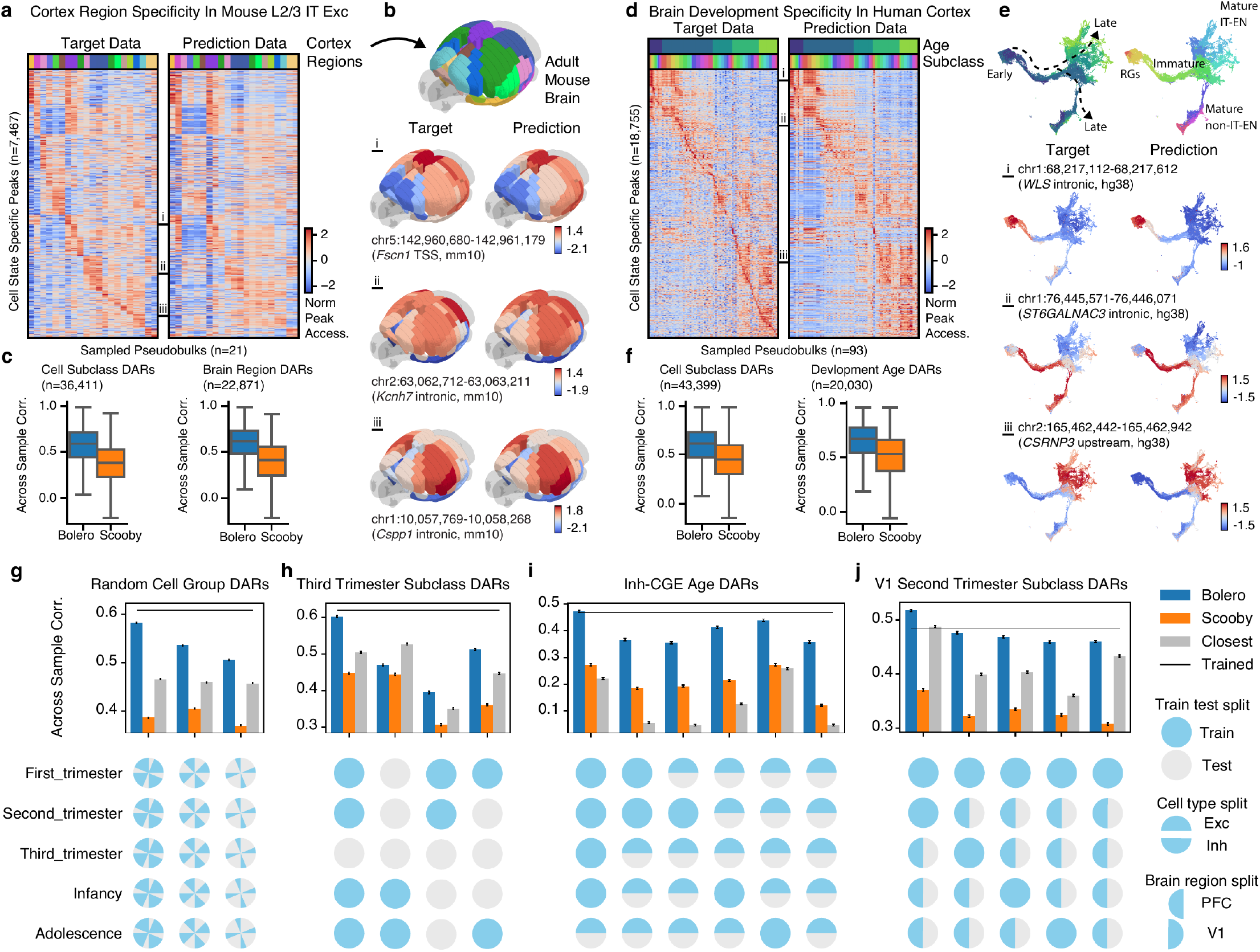
Bolero predicts spatial and temporal regulation, including in held-out cell states. **a,** Heatmaps of measured (left) and predicted (right) accessibility in “L2/3 IT neurons” across 21 brain regions for DARs between brain regions. Accessibility was quantified in counts per million (CPM) after log1p and double z-scoring. **b,** Common coordinate framework map coloured by brain regions (top), or measured (left) and predicted (right) accessibility for example peaks. **c,** Pearson correlation between measured and predicted accessibility across subclass x brain region pseudobulks over DARs between subclasses (n = 36,411) or brain regions (n = 22,871). **d,** Heatmaps of measured (left) and predicted (right) accessibility across human developmental stages and excitatory subclasses. **e,** UMAP of excitatory lineage coloured by pseudotime (top left), subclass (top right), or measured (left) and predicted (right) accessibility for example peaks. **f,** Pearson correlation between measured and predicted accessibility across subclass x development stage pseudobulks over DARs between subclasses (n = 43,399) or developmental stages (n = 20,030). **g–j,** Pearson correlation between measured and predicted accessibility across randomly held-out cell groups (three held-out fractions, g; 52 pseudobulks, 32,870 DARs), third trimester subclasses (h; 25 pseudobulks, 10,381 DARs), Inh-CGE (caudal ganglionic eminence) lineage age groups (i; 21 pseudobulks, 4,573 DARs), or V1 second trimester subclasses (j; 21 pseudobulks, 17,195 DARs) on corresponding DARs (panel title), using Bolero or Scooby with correct condition and embedding, or Bolero with closest condition and embedding among training pseudobulks. Bolero performances when trained on all cell groups are shown as black lines. Error bars denote s.e.m. across peaks. The train-test split map for each experiment is shown at the bottom.

For the temporal axis, we used a human cortex single-cell multiomic atlas covering five developmental stages from the first trimester to adolescence and two cortical regions, the prefrontal cortex (PFC) and the primary visual cortex (V1)^3^. Bolero’s predictions tracked both the subclass and the developmental-age axes across 18,755 cell-state-specific peaks and 93 pseudobulks (subclass-by-age), recovering the orderly progression from early radial glia to mature IT and non-IT excitatory neurons (Fig. 2d, e). Predicted accessibility at developmental cCREs near *WLS*, *ST6GALNAC3*, and *CSRNP3* tracked the measured developmental-stage signal (Fig. 2e). On cell-subclass DARs and developmental-age DARs, Bolero’s median across-sample correlation was 0.61 and 0.68, compared with 0.45 and 0.53 for Scooby, an improvement of about 30% (Fig. 2f).

A more demanding test of the efficacy of Bolero is generalisation to cell states absent from the training data. We retrained Bolero with four progressively challenging held-out splits: random cell groups; random developmental time points; all inhibitory neurons at given time points; and all visual cortex cells at given time points (Extended Data Fig. 4a). We then benchmarked the resulting predictions against two reference points. “Closest” runs the same Bolero model on the held-out genomic regions but replaces the cell-state embedding and metadata-conditioning input with those of the nearest training pseudobulk; this isolates whether the actual held-out embedding carries information beyond the closest training cell-states. “Trained” is an upper bound obtained from a model trained directly on the held-out cells. Across the four splits, Bolero with the held-out embedding exceeded “Closest” in nearly every held-out case (17 of 18 featured cases) and approached “Trained” (Fig. 2g–j, Extended Data Fig. 4b,c). The same result held when the conditioning embedding was computed from RNA alone (scVI, scANVI) rather than from the multiome (MultiVI), so it does not depend on ATAC information reaching the model through the embedding (Extended Data Fig. 4b). This indicates that the embedding carries continuous information about cell state, beyond the identity of the nearest training sample, on which sequence regulation can be evaluated for cell types, brain regions, or developmental stages absent from training.

### Cell-state-resolved variant-effect prediction connects sequence to disease

One of the most consequential applications of sequence-to-function models is to interpret non-coding genetic variation: predicting which variants alter regulation, and in which cell types. The clinical and population-genetics utility of these predictions has been limited by the same cell-state-resolution gap that motivates Bolero, since bulk-tissue effect estimates obscure the cell types in which a variant has an impact. We tested Bolero’s cell-state-conditioned architecture in three settings: per-tissue Quantitative Trait Locus (QTL) benchmarks; population-scale Genome-Wide Association Study (GWAS) heritability partitioning; and de novo mutation interpretation in autism.

Bolero performs in silico substitution to predict the cell-state-specific change in chromatin accessibility and transcript abundance for any candidate variant (Fig. 3a). QTL studies associate genotypes with molecular readouts (e.g. chromatin accessibility, gene expression) across individuals, and fine-mapping then identifies the variants that are most likely causal. We first used these datasets to test Bolero’s performance on variant effect predictions on two axes. The first is whether the predicted effect size agrees with the measured effect at fine-mapped variants. The second is whether Bolero ranks fine-mapped QTLs above the far larger set of tested variants with no detectable effect. For chromatin accessibility QTL (caQTL) on adult cortical neurons and glia^26^, predicted and measured effect sizes agreed (neurons, Fig. 3b; glia, Extended Data Fig. 5a). At ranking fine-mapped caQTLs, Bolero matched the strongest task-specific baseline Scooby and exceeded ChromBPNet (neurons, Fig. 3c; glia, Extended Data Fig. 5b). For expression QTL (eQTL) on the OneK1K single-cell benchmark spanning 19 blood cell types^27^ and on whole-blood GTEx fine-mapped variants, predicted and measured effect sizes again agreed (Fig. 3b, Extended Data Fig. 5c). At ranking fine-mapped eQTLs, Bolero matched Decima and exceeded Borzoi and a baseline built on distance to the transcription start site (TSS; Fig. 3c). Bolero therefore achieves per-cell-type accuracy comparable to the strongest bespoke models.

**Figure 3.**
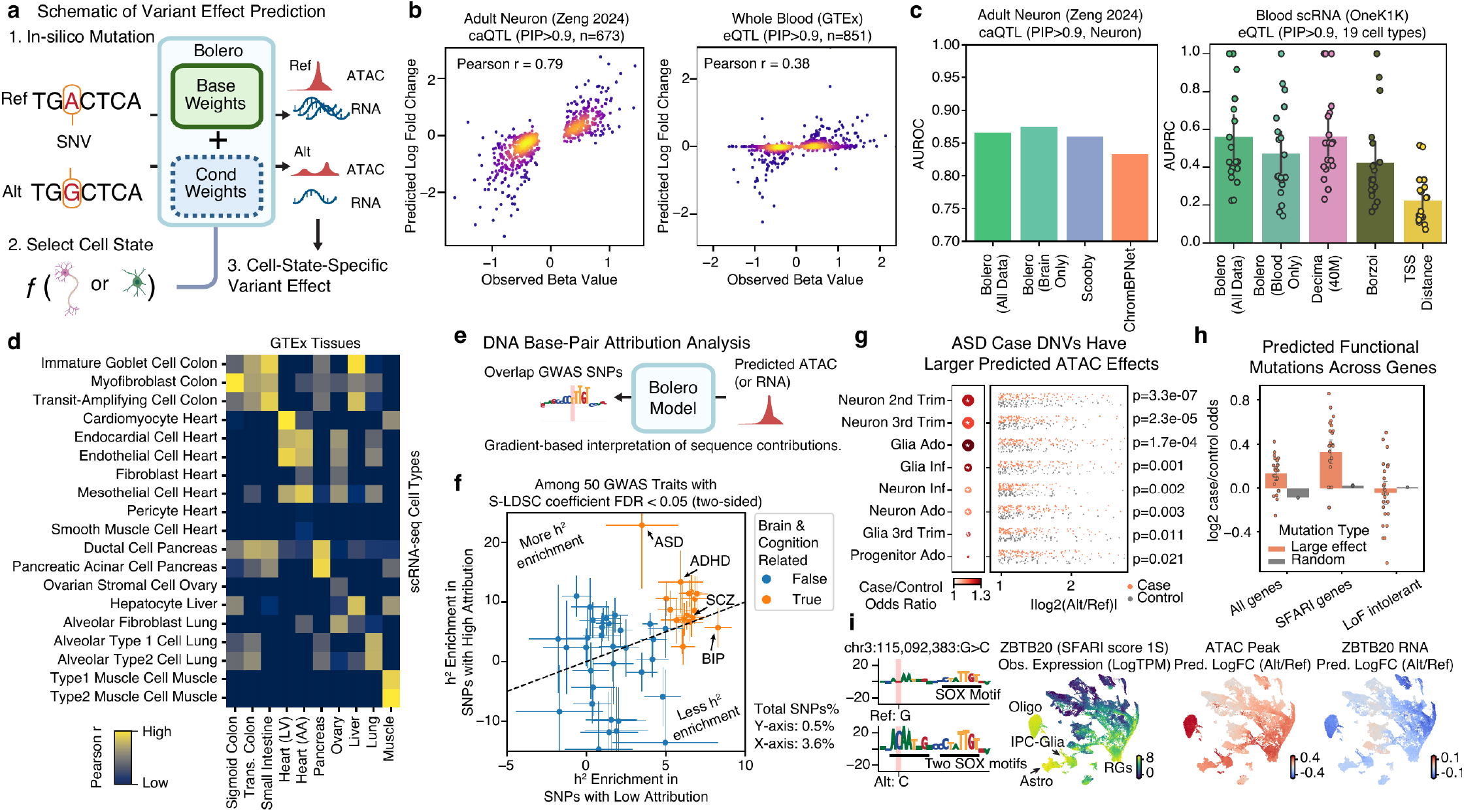
Cell-state-resolved variant-effect prediction connects sequence to disease. **a,** Schematic of variant-effect prediction. **b,** Observed versus predicted effects for adult-neuron caQTLs (Zeng 2024, posterior inclusion probability (PIP) > 0.5, n = 673; Pearson r = 0.79) and whole-blood eQTLs (GTEx, PIP > 0.9, n = 851; Pearson r = 0.38). **c,** Average precision (AUPRC) for prioritising fine-mapped blood eQTLs across 19 immune cell types (OneK1K, PIP > 0.9; n = 957 positive and 21,617 negative variants; error bars, 95% CI) and AUROC for prioritising fine-mapped adult-neuron caQTLs (Zeng 2024, PIP > 0.5; n = 673 positive and 1,094 negative variants), for different methods. **d,** Pearson correlation between Bolero’s predicted eQTL effects in each ENCODE Human Atlas cell type (rows) and the observed effects of each GTEx tissue’s eQTLs (columns). **e,** Schematic of the gradient-based per-base attribution workflow used to interpret sequence contributions and overlap them with GWAS SNPs. **f,** Heritability enrichment (S-LDSC, two-sided) for SNPs with high attribution (0.5% of SNPs) versus low attribution (3.6%), among 50 GWAS traits passing coefficient FDR < 0.05; brain and cognition traits highlighted; error bars, standard error. **g,** Predicted ATAC effect size of autism case versus control de novo variants per cell class and developmental stage, with the case-versus-control odds ratio (n = 127,137 variants; 65,146 proband and 61,991 sibling variants; asterisks denote significance). **h,** Case-versus-control odds ratio of non-coding de novo variants in three gene categories, computed for large-effect variants in each cell class and developmental stage (bars, mean; dots, class x stage groups; error bars, 95% CI) and, as a baseline without selection on predicted effect, for all de novo variants (Random, the case-to-control mutation ratio; one value per category). **i,** DNA sequence across the window centred on the variant with letter height representing reference-allele (top) and alternate-allele (bottom) per-base attribution with the variant highlighted (left); and UMAPs of HumanBrainDev cells (n = 228,630) coloured by observed ZBTB20 expression, predicted ATAC-peak log fold-change (Alt/Ref), and predicted ZBTB20 RNA log fold-change (right) at metacell resolution.

Bolero predicted almost no effect for 76.5% of fine-mapped eQTLs, raising the question of whether these misses reflect a limitation of the model or a property of the variants themselves. Stratifying eQTLs into predictable and unpredictable subsets, we found that predictable variants were enriched close to the TSS, in promoters and 5’UTRs, in chromatin carrying marks of active regulation, and, modestly, among variants predicted to disrupt splicing. The unpredictable set shifted toward distal intergenic regions farther from the TSS, but remained heterogeneous and could not be assigned cleanly to a single mechanism (Extended Data Fig. 5d–i). The distal shift is consistent with limited sensitivity to long-range regulatory variation, a general property of current sequence-to-function models^28^. The stratification identifies which variants Bolero can be trusted on before downstream interpretation.

Cell-state resolution emerges most clearly when bulk QTL effects are decomposed into the cell types from which they originate. We collected 29,248 fine-mapped eQTLs from 50 tissues in GTEx, and predicted their effects on the expression of their linked genes across 38 cell types in the ENCODE scRNA-seq atlas. Using the dominant cell types in a particular tissue generally yields predictions that are closer to the measured effects. For example, heart eQTLs aligned to cardiomyocytes and to endocardial, endothelial and mesothelial subsets, lung eQTLs to “alveolar Type 1/2 cells”, and pancreas eQTLs to ductal and acinar cells (Fig. 3d). The same decomposition resolved individual whole-blood eQTLs into their constituent peripheral-blood mononuclear-cell (PBMC) types, agreeing with the effects independently fine-mapped to those cell types in the OneK1K cohort (Extended Data Fig. 6a,b). When resolved further to per-base attribution, the reference-minus-alternate change recovered the cell type expected from the source dataset of the QTL and localised on the motif sequence (Extended Data Fig. 6c,d). The two examples were “CD16+ NK cells” for an immune eQTL at chr5:1,104,823 C>T, and “L2/3 IT excitatory neurons” for a brain eQTL at chr2:176,628,649 G>A.

Beyond per-variant scoring, Bolero’s gradient-based attribution scores assign every base of every cell-type-specific open chromatin peak a contribution to predicted accessibility (Fig. 3e). Within the union of HumanBrainDev dataset neuronal open-chromatin peaks, we partitioned the peak set into high-attribution sub-regions and the remaining peak interior. We ran stratified Linkage Disequilibrium (LD)-score regression (S-LDSC) on top of the standard baseline LD model^29,30^. Among 50 GWAS traits with significant Bolero S-LDSC coefficients (FDR < 0.05), heritability enrichment was systematically higher in the high-attribution sub-regions. Enrichment was highest for brain and cognition phenotypes, reaching ∼20-fold for Autism Spectrum Disorder (ASD) and exceeding 7-fold for schizophrenia and Attention-Deficit/Hyperactivity Disorder (ADHD) (Fig. 3f). The high-attribution fraction of the open neuronal genome carries a disproportionate share of polygenic risk for psychiatric disease, identifying a small subset of the genome where Bolero’s sequence model converges with the heritability signal of population-scale GWAS.

The same predictions resolve disease genetics down to individual non-coding de novo variants (DNVs). Across a large autism case-control cohort^31^, case DNVs produced larger predicted effects on peak accessibility than control DNVs, especially in early neuronal developmental stages (Fig. 3g). To ask whether that signal tracks disease relevance, we compared the case-versus-control odds of large-effect variants across three gene sets: all genes, autism-risk genes^32^, and loss-of-function-intolerant genes^33^. The last of these are essential but not specifically autism-associated and serve here as a control. Enrichment was modest across all genes, strongest at autism-risk genes, and absent at loss-of-function-intolerant genes (Fig. 3h). The predicted effects therefore track autism relevance rather than gene essentiality, indicating that Bolero captures functional variant effects at disease-relevant genes. As an example, a non-coding de novo G>C substitution in *ZBTB20* (SFARI risk score 1S; chr3:115,092,383) creates a SOX-family binding motif on each strand that Bolero predicts increases the peak’s chromatin accessibility and decreases *ZBTB20* expression in oligodendrocyte and astrocyte lineages (Fig. 3i). This is a cell-state-specific, direction-resolved prediction that bulk-tissue models cannot resolve.

Across QTL benchmarks, GWAS heritability, and DNV analysis, Bolero’s predictions track disease genetics from genome-wide partitioning down to individual variants, with cell-state resolution that makes the predictions interpretable.

### AP-1-conditioned Bolero predicts and designs cell-type-specific activity-regulated cCREs

Neuronal activity induces a stereotyped immediate-early gene (IEG) program dominated by AP-1 family transcription factors, yet the enhancers activated downstream of this shared induction signal are highly cell-type-specific^34,35^. The regulatory logic that translates a shared induction signal into divergent cell-type programs is unresolved, in large part because activity-regulated chromatin datasets remain available for only a limited number of paradigms and cell states. Yet the underlying cis-regulatory grammar must be inferred across many unmeasured combinations of cell type, developmental stage, species, and stimulus context. We asked whether Bolero could predict AP-1 activity responses across cell states after training on the multi-atlas datasets. Specifically, we added a per-cell AP-1 chromVAR activity score^36^ as an additional conditioning variable, which quantifies the chromatin accessibility of AP-1 motifs in single cells, alongside the cell-state embedding. We fine-tuned the model jointly with this score to obtain Bolero-AP1-Score, a model that predicts AP-1-modulated, cell-state-specific chromatin accessibility from DNA sequence (Fig. 4a).

**Figure 4.**
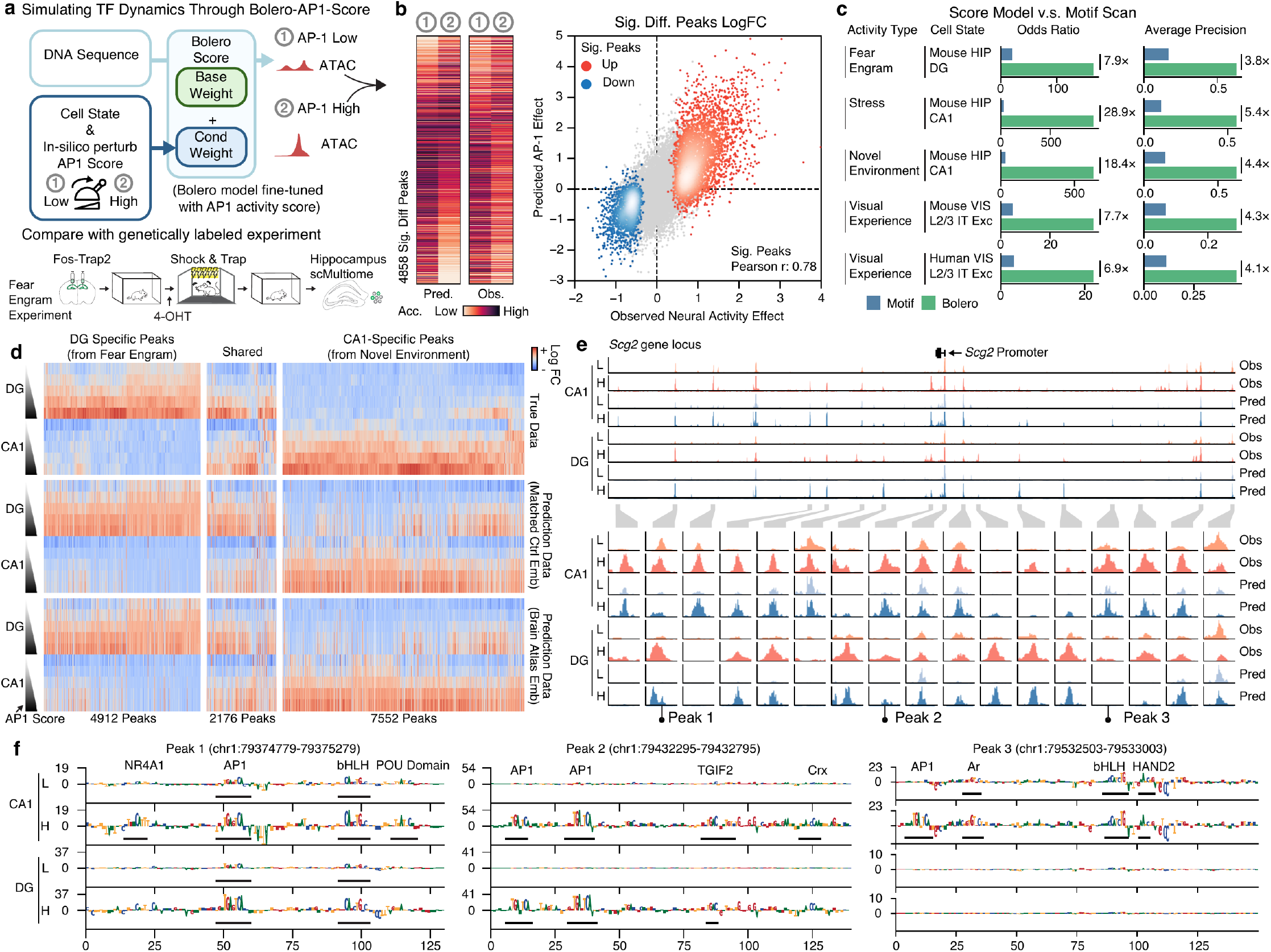
Bolero-AP1-Score predicts stimulus-dependent accessibility. **a,** Top, schematic of Bolero-AP1-Score: both cell state and AP-1 score condition the model to predict accessibility from DNA sequence. Bottom, schematic of the genetically labelled fear-engram experiment used for comparison: Fos-Trap2 labelling with 4-OHT during fear conditioning, followed by hippocampal single-cell multiome profiling. **b,** Heatmaps of predicted (AP-1-low versus AP-1-high) and measured (Trap− versus Trap+) accessibility at significant differential peaks in mouse hippocampal DG in a fear-engram paradigm (n = 4,858, left); and predicted versus observed AP-1 effect across peaks (n = 36,469; 3,293 up, 1,565 down) coloured by density, with significantly up in red, down in blue, or non-significant in grey (right). **c,** Odds ratio for the predicted direction (up versus down) of significant differential peaks, and average precision for prioritising the significantly up-regulated peaks against all others, of Bolero-AP1-Score (green) versus a motif scan (blue) across five paradigms, with fold improvement annotated between the paired bars. **d,** Measured and predicted accessibility (colour) at DG-specific (n = 4,912), shared (n = 2,176), and CA1-specific (n = 7,552) peaks (column blocks), with true data and predictions using a matched-control embedding or a brain-atlas embedding (row blocks), each split into DG and CA1 cell states with rows ordered by AP-1 score. Values are z-scored per peak; peaks within each column block are ordered by Ward hierarchical clustering. **e,** Measured (orange) and predicted (blue) accessibility tracks across the Scg2 locus in DG and CA1 at AP-1-low and AP-1-high (top), with zoomed profiles at individual peaks (bottom). **f,** DNA sequence at three example Scg2-locus peaks with letter height representing per-base attribution, each for CA1 and DG at AP-1-low and AP-1-high, with de novo seqlets annotated to TF motifs.

The key premise of this design is that AP-1 chromVAR provides a quantitative proxy for the AP-1-associated component of neuronal activity. We first evaluated this premise across five independent stimulation or experience paradigms (Supplementary Table 4). These included fear conditioning in mouse hippocampal dentate-gyrus (DG) engram cells, stress and novel-environment exposure in mouse CA1^37,38^, visual experience in mouse visual cortex (VIS) L2/3 IT excitatory neurons^4^ and in human V1 L2/3 IT excitatory neurons^3^. In three of the five paradigms (fear conditioning, novel environment and human visual experience), AP-1 showed the largest chromVAR activity-score change of any transcription-factor motif between stimulated and control conditions, and the second largest in stress and mouse visual experience (Extended Data Fig. 7a). This means that AP-1 is the most reproducible motif-level signature of activity across brain regions, stimuli, and species. AP-1 activity increased along the visual-cortex developmental trajectory in both mouse and human L2/3 IT neurons (Extended Data Fig. 7b). During fear memory, it separated engram-labelled FOS-Trap+ cells from FOS-Trap− cells in a cell-type-resolved manner across hippocampal and glial populations (Extended Data Fig. 7c). These observations support the use of AP-1 chromVAR as a continuous conditioning variable for modelling activity-associated chromatin responses.

We next tested whether Bolero-AP1-Score could predict activity-induced chromatin changes in behaviour-paired paradigms. We trained Bolero-AP1-Score on Bolero-10M while holding out all the stimulus data from the mouse behaviour experiments. In the fear-engram dataset, the predicted AP-1 effect closely tracked the measured activity-induced accessibility change at DARs (Pearson r = 0.78; Fig. 4b). Across all five paradigms, Bolero-AP1-Score predictions tracked the direction and magnitude of observed accessibility changes at individual peaks (Extended Data Fig. 7d). In contrast, AP-1 motifs were enriched among positively regulated peaks but were substantially less predictive of individual peak responses (Extended Data Fig. 7d). Consistently, Bolero-AP1-Score substantially outperformed a motif-scan baseline in prioritizing activity-induced peaks, both by odds ratio and by average precision, with the odds ratio improving up to 29-fold across the evaluated paradigms (Fig. 4c). Thus, activity-associated chromatin responses are encoded in cis at individual regulatory elements and can be recovered by a model that never saw activity-induced chromatin during training.

AP-1 induction is a common response to neuronal activity, but its consequences are cell-type-specific, because AP-1 binds different cCREs in different cells. We asked whether Bolero-AP1-Score could predict these cell-type-specific cCRE responses. Across hippocampal activity-regulated peaks, including DG-specific fear-engram peaks and shared or CA1-specific novel-environment peaks, Bolero-AP1-Score reproduced the expected DG-versus-CA1 responses. This was true both when the model was conditioned on cell-state embeddings computed only from non-stimulated control cells from the same experiment and when it was conditioned on broader brain-atlas embeddings (Fig. 4d). Cell identity and AP-1 activity are therefore represented as separable inputs: the embedding specifies the cellular regulatory context, while the AP-1 score specifies the activity-associated axis. At a cCRE near the canonical activity-regulated gene *Scg2*, the predicted DG/CA1 × AP-1-low/high accessibility pattern matched the measured pattern (Fig. 4e). Per-base attribution at three representative peaks further highlighted sequence features consistent with cooperative motif logic, including AP-1 sites paired with NR4A1, POU-domain, bHLH, TGIF2, Crx, HAND2, or AR motifs in a peak- and cell-type-dependent manner (Fig. 4f). These results suggest that a shared AP-1 signal is interpreted through local combinations of cooperating motifs to generate divergent cell-type-specific cCRE outputs.

If Bolero-AP1-Score has learned this cis-regulatory grammar, the same model should be able not only to read activity-regulated cCREs but also to design sequence changes that redirect their predicted activity. We tested this with directed sequence evolution. Starting from a *Scg2* cCRE (Peak 3) that shows AP-1-dependent accessibility in CA1 while remaining largely closed in DG, we sampled point mutations and scored each variant against two target cell types × two AP-1-states accessibility patterns. We then iteratively retained mutants that moved toward the desired target (Extended Data Fig. 8a). Designed Peak 3 variants reached each requested cell-type × AP-1-state pattern with sparse sequence changes, corresponding to mutation rates of ∼0.33–2% (Extended Data Fig. 8b-e). The prioritised mutations exposed an interpretable design logic. To suppress the AP-1 response in CA1, the model preferentially removed AP-1 motifs; to more completely suppress CA1 accessibility, it removed additional bHLH motifs (Extended Data Fig. 8f). Conversely, to activate basal accessibility in DG, the model introduced bHLH and NF-I motifs, and to introduce an AP-1 response in DG, it further added AP-1 motifs (Extended Data Fig. 8g). Thus, the same cooperative motif logic inferred from natural cCREs can be used for in silico sequence design, supporting Bolero-AP1-Score as an interpretive framework for cell-type-specific activity-regulated cis-regulatory logic.

### Bolero-AP1-Score reconstructs the context-dependent cooperative grammar of AP-1 responses

AP-1 has long been viewed as a broadly inducible transcription-factor hub whose genomic specificity is imposed by cooperative interactions with lineage-determining factors, chromatin remodelers, and local sequence context^39–42^. There are ∼600,000 potential AP-1 sites in the genome, and each cell type is predicted to use its own unique repertoire of sites that then determine the cell response to extracellular stimuli, including proliferation, differentiation, oncogenic transformation, and long-term memory formation, maintenance, and retrieval^34^. Having established that Bolero-AP1-Score can recover activity-regulated cCREs in measured neuronal paradigms, we next asked whether the same model could expose the sequence logic by which AP-1 responses are specified across the human body (Fig. 5a). We predicted chromatin accessibility with AP-1-high versus AP-1-low across 164 human-body cell pseudobulks and compared the genome-wide AP-1 response profiles between cell states. These predicted response profiles were not uniform across all AP-1-containing regulatory elements, but instead clustered by major cell-class groups, including brain, mesenchymal, myeloid, lymphoid, epithelial, glandular/endocrine, and endothelial populations (Fig. 5a,b). This suggests that Bolero-AP1-Score has learned a context-dependent regulatory grammar rather than a simple AP-1 motif score.

**Figure 5.**
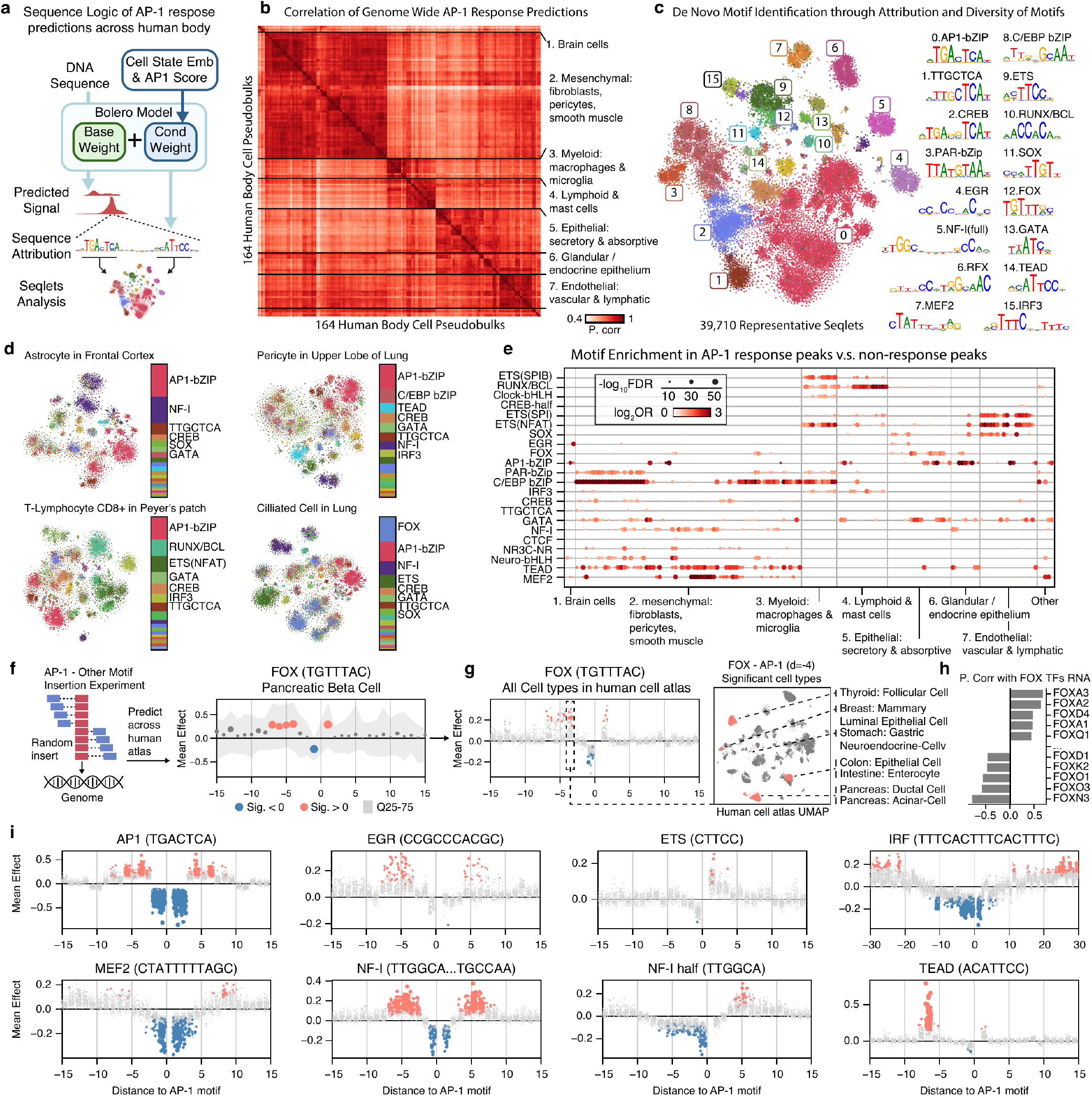
Bolero-AP1-Score reconstructs the context-dependent cooperative grammar of AP-1 responses. **a,** Schematic of seqlet analysis. **b,** Genome-wide Pearson correlation of the predicted AP-1 response between human-body cell pseudobulks (n = 164, rows and columns), Ward-clustered into seven cell-class groups. **c,** t-SNE of representative seqlets identified across all 1,204 training pseudobulks based on their per-base attribution, coloured by de novo motif cluster with its consensus logo and TomTom annotation (16 selected clusters labelled). **d,** t-SNE of seqlets based on attribution for four example cell types (each embedded separately), coloured by the motif families recovered in each. **e,** Enrichment of each de novo motif family (rows) in AP-1-response versus non-response peaks across pseudobulks (columns, n = 153). **f,** Left, schematic of the motif-insertion assay. Right, mean predicted AP-1 effect (log2 fold-change) versus the signed distance between the inserted FOX and AP-1 motifs in pancreatic beta cells (gap in bp between the motif edges; negative values denote a partner motif on the reverse strand); dots coloured by the sign of a significant effect (one-sample t-test versus zero, BH-FDR < 0.05) or grey (not significant); ribbon, interquartile range across constructs. **g,** Mean FOX–AP-1 effect versus distance pooled over all 164 pseudobulks (left; each dot a pseudobulk × spacing, the d = −4 spacing boxed), and a human cell-atlas UMAP (right) with cell types showing a significant effect at d = −4 highlighted. **h,** Pearson correlation across cell subclasses (n = 41) between the FOX–AP-1 effect (per-subclass t-statistic at d = −4) and RNA expression of FOX-family transcription factors, showing the five most positively and five most negatively correlated FOX genes. **i,** Mean effect versus signed distance to the AP-1 motif for additional partner motifs (each dot a pseudobulk × spacing, coloured as in f).

To identify the sequence features underlying this structure, we computed AP-1-high versus AP-1-low attributions and applied TF-MoDISco^43^ to discover de novo seqlets (short, high-attribution stretches of sequence) from predicted AP-1-responsive regions. This yielded 39,710 representative seqlets that resolved an AP1-bZIP core together with a broad repertoire of candidate cooperator motifs (Fig. 5c). The resulting motif catalogue recapitulated known AP-1 partner classes, including NFAT/ETS-like and IRF composite logic in immune cells^44,45^, PU.1/ETS and C/EBP-associated logic in myeloid contexts^46^, FOXA-associated epithelial programs^47,48^, and TEAD-associated vascular or mesenchymal programs^49^. Consistent with this interpretation, different cell types used distinct motif repertoires around the shared AP-1 axis: “lung ciliated cells” were enriched for FOX, NF-I, and ETS-like motifs; “lung pericytes” for TEAD, C/EBP, and NF-I motifs; “Peyer’s patch CD8+ T cells” for RUNX/BCL and ETS/NFAT-like motifs; and “frontal-cortex astrocytes” for NF-I, SOX, and CREB-like motifs (Fig. 5d). Enrichment analysis further showed that AP-1-response peaks were enriched not only for AP1-bZIP motifs, but also for cell-class-associated cooperators relative to non-response peaks (Fig. 5e). These results place known AP-1 cooperative partners into a unified atlas-scale map, in which each cell class uses a different cooperator vocabulary to interpret a shared AP-1 signal.

Because motif enrichment alone cannot distinguish passive co-occurrence from cooperative syntax, we next converted the motif catalogue into a perturbational assay of motif identity and spacing. We inserted AP-1 and candidate cooperator motifs into controlled sequence backgrounds, varied the signed distance between the motif pair, and predicted the resulting AP-1 effect across the human atlas (Fig. 5f). We first focused on FOX as an illustrative example. Prior work showed that AP-1 can promote FOXA1 binding at nearby sites, with cooperativity strongest at close motif proximity and dependent on cellular AP-1 activity^47^. In Bolero-AP1-Score, inserting FOX motifs beside AP-1 motifs similarly revealed a spacing-dependent effect (Fig. 5g). This effect was not globally shared across all cell types but was concentrated in epithelial and glandular/endocrine states, consistent with the idea that motif syntax is interpreted through the TF repertoire available in each cellular context (Fig. 5g). Accordingly, the strength of FOX–AP-1 cooperativity correlated with FOX-family RNA expression across cell types, nominating candidate FOX genes (e.g. FOXAs) that could mediate the response (Fig. 5h). Beyond FOX, many additional cooperators showed distance-dependent modulation of the AP-1 response, including EGR, ETS, MEF2, NF-I, TEAD, and IRF motifs, each with its own characteristic spacing preference and cell-class specificity (Fig. 5i; Extended Data Fig. 9). Thus, inserting motif pairs into controlled backgrounds transforms the attribution-derived cooperator catalogue into a quantitative syntax model, linking AP-1 response strength to motif identity, local spacing, and cell-state-specific TF availability.

Finally, we used AlphaFold3^50^ as an orthogonal, hypothesis-generating analysis to ask whether the model-inferred spacing effects could reflect DNA-mediated TF co-binding. As a representative example, AP-1–TEAD was selected because the insertion assay showed a particularly sharp spacing- and orientation-dependent effect, and because AP-1–TEAD composite elements have been reported in vascular and hemogenic regulatory programs^40,49,51^. AlphaFold3 models of AP-1 with TEAD1/YAP1/TAZ on DNA showed spacing-dependent regions of confidently predicted contact between FOS or JUN and the cooperative factors (low predicted aligned error, PAE) near the spacings predicted by the insertion assay (Extended Data Fig. 10a-c). We extended this analysis across the broader set of candidate AP-1 cooperators. Confident AP-1–cooperator interaction segments were more evident in DNA-containing structure models than in no-DNA controls, consistent with DNA-mediated co-binding rather than stable protein–protein association alone (Extended Data Fig. 10d). Together, these analyses extend known AP-1 composite-element logic to an atlas-scale predictive grammar in which AP-1 responsiveness is specified by motif identity, local spacing, and cell-state-specific cooperator context.

### Bolero generalises cell-state-specific and AP-1 predictions across mammals

Genome assemblies exist for hundreds of mammals^52,53^, but matched single-cell chromatin and transcriptomic atlases cover only a handful of species due to cost and labour limitations. Comparisons of gene regulation between species have been confined to the few species anyone has profiled. More importantly, a synchronous stimulus can be delivered to mice or non-human primates, but the inducible fraction of the human regulatory genome has never been mapped directly. That fraction could harbour what distinguishes human brain function, and be involved in complex diseases. Bolero has internalised cell-state-specific regulation in one or two species and could, in principle, be queried directly from the genome of any other species, turning a genome browser into a regulatory-function browser (Fig. 6a).

**Figure 6.**
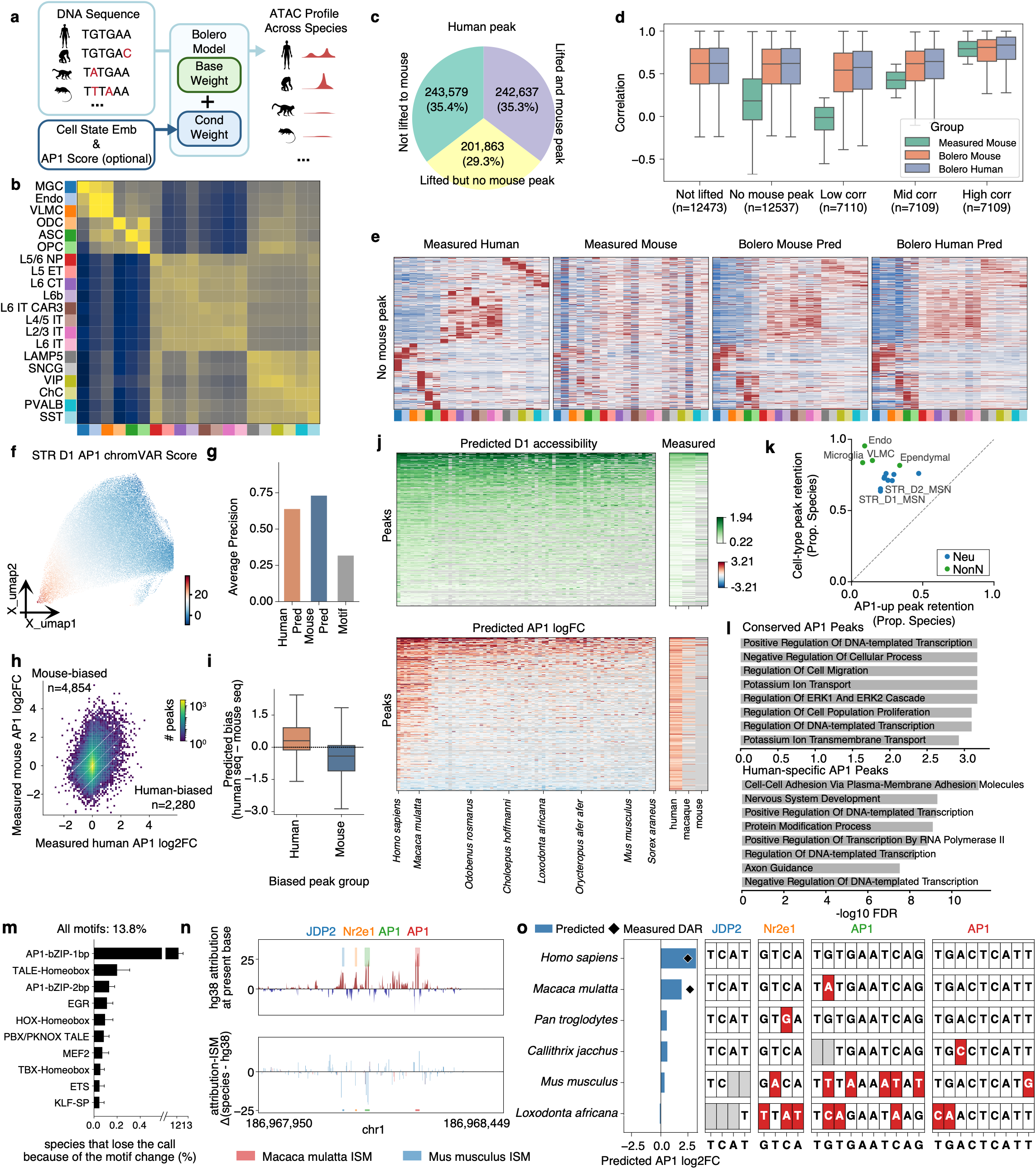
Bolero generalises cell-state-specific and AP-1 prediction across mammals. **a,** Schematic of cross-species prediction. **b,** Across-subclass correlation matrix over the 20 shared cortical subclasses between measured human accessibility (rows) and prediction from Bolero mouse model using human sequence (columns). **c,** Human-to-mouse peak-liftover outcome as a pie chart. **d,** Per-peak Pearson correlation across the 20 subclasses within each conservation group between measured human accessibility and measured mouse accessibility at orthologous peaks, or the prediction from Bolero mouse or human models with human sequence (sample sizes labelled in the plot). **e,** Heatmap of measured accessibility in human (Measured Human), in mouse at liftover coordinates (Measured Mouse), or predicted with human sequence by Bolero model trained across mouse (Bolero Mouse Pred) or human (Bolero Human Pred) datasets, in human DARs that lift over to mouse but fall in no mouse peak (the “No mouse peak” group of (c) and (d); columns, colour shared with (b)). Values are z-scored per peak across subclasses. **f,** UMAP of human striatal D1 medium spiny neurons coloured by AP-1 (MA0478.2) chromVAR score. **g,** Average precision for classifying mouse AP-1-up peaks (7,032 of 76,129) from all D1 peaks with Bolero model trained with human (Human Pred) or mouse (Mouse Pred) data, and a motif scan (Motif). **h,** Measured human versus measured mouse AP-1 log2 fold-change at orthologous peaks as a 2D density (colour, peaks per bin, log scale), with an identity line and offset guide lines demarcating mouse-biased (n = 4,854) and human-biased (n = 2,280) peak groups. **i,** Predicted AP-1 log fold-change from human minus mouse sequence for the human-biased and mouse-biased peak groups. **j,** Cross-mammalian heatmaps of predicted D1 accessibility (top) and predicted AP-1 log fold-change (bottom) at alignable D1 peaks (sampled rows), columns being human and the 74 non-human genomes into which at least 500,000 human peaks were lifted, ordered by sequence divergence from human, with a narrower measured heatmap across human, macaque, and mouse (right). Peaks that could not be lifted over are coloured grey. **k,** Fraction of species retaining the predicted signal for AP-1-inducible (x) versus cell-type (y) peaks across major cell types (dot, n = 13; retention over 83 species), coloured by neuronal vs non-neuronal. **l,** Gene Ontology biological-process terms enriched for genes associated with conserved (top) and human-specific (bottom) AP-1 peaks, showing the eight most significant terms per set as -log10 FDR; dashed line, FDR = 0.05. **m,** Percentage of species predicted to lose the AP-1 response owing to changes at attributed motif sites, showing the ten largest motif contributions across 8,780 AP-1-responsive D1 peaks (broken x-axis). Error bars are 95 percent bootstrap intervals over peaks, from 1,000 draws. **n,** Per-base attribution of an example D1 peak near PLA2G4A/PTGS2, with JDP2, Nr2e1, and two AP-1 seqlets highlighted (top), and attribution-based in silico mutagenesis tracks (species minus human) for macaque and mouse (bottom). **o,** Per-species predicted (bars) and measured (diamonds, human and macaque only) AP-1 log2 fold-change at that enhancer across six species (left); and each of the four seqlets’ (JDP2, Nr2e1 and the two AP-1) sequences in the six species with mismatches to human highlighted (right). For all box plots, the centre line denotes the median; box limits denote first and third quartiles; and whiskers denote 1.5 × the interquartile range.

We first tested whether a model trained in one species can recover the cell-type structure of another. A version of Bolero trained on mouse data predicted human cortical chromatin best for the matched human cell type, indicating that cell-type-specific open chromatin can be recovered from another species’ sequence (Fig. 6b). We next examined how predictions behave across different peak types. About 35% of human cortical peaks had no syntenic mouse counterpart, and another 29% could liftOver to the mouse genome but fell within inactive chromatin (Fig. 6c), restricting traditional cross-species peak comparison to the remaining ∼35%. Bolero predicted all human peaks directly from a mouse data-trained model, capturing cell-type-specific signals at human-only peaks as well as at shared peaks whose cell-type specificity differs between species (Fig. 6d, e, Extended Data Fig. 11a). Predictive performance for human open chromatin decreased only modestly with the evolutionary distance of the training species, whereas the cross-species similarity of measured signals declined far more steeply (Extended Data Fig. 11b). These results suggest that Bolero generalizes well across species by capturing regulatory signatures in DNA sequence rather than memorizing species-specific peak repertoires.

Beyond cell-type-specific chromatin prediction, we turned to activity-dependent regulation, where the gap between species is starkest. We focused on D1 medium spiny neurons (D1 MSNs) of the basal ganglia, whose AP-1 activity varies along an internal dorsal-ventral gradient mirroring the anatomical axis running from the caudate/putamen through the nucleus accumbens to the olfactory tubercle^54–56^. AP-1 activity defines a dominant continuous axis of D1 chromatin accessibility variation (Fig. 6f). Similar to the mouse hippocampus experiment, we identified 19,725 (7,032) DARs upregulated in AP-1 high versus low cells in humans (mice) as AP-1 responsive peaks. Trained on human data alone, the AP-1-conditioned model predicted AP-1-induced chromatin accessibility changes of D1 cCREs in mice, from the mouse genome, more accurately than a motif-scan baseline (Fig. 6g). To test whether species-specific AP-1 responses are also captured, we predicted AP-1 responses for the same orthologous peaks using both human and mouse sequences and compared the predictions. Bolero predicted higher AP-1 responses from human than from mouse sequence at human-biased peaks, and the reverse at mouse-biased peaks (Fig. 6h,i). The same pattern held between human and macaque sequences (Extended Data Fig. 11c). These results suggest that species-specific tuning of the AP-1 response is partially encoded in cis and recoverable from sequence alone.

The accuracy of these cross-species predictions enabled us to extend comparative analysis from sequence conservation to functional conservation. We predicted accessibility and AP-1 responses across 84 Zoonomia mammals (human and 83 other species) at every alignable D1 MSN peak (Fig. 6j). AP-1-responsive peaks show sequence conservation comparable to other D1 peaks (Extended Data Fig. 11d), but their functional conservation is substantially weaker than that of D1 peaks overall (Fig. 6j). For each of the 13 major basal-ganglia cell types examined, we compared the number of species retaining a peak’s chromatin accessibility against retaining its AP-1-inducible signal. The chromatin accessibility was retained in more species than AP-1 responses in all 13 cell types (Fig. 6k). The activity-inducible layer of basal-ganglia cCREs is therefore less conserved than the cell-identity layer built on the same elements. Among the 10,385 AP-1 responsive peaks in humans, we identified 1,178 peaks showing conserved responses across species, 3,004 showing primate-specific responses, and 6,203 with human-specific responses. Conserved AP-1 peaks were enriched for core transcriptional and ERK-related regulatory programs. In contrast, human-specific AP-1 peaks showed stronger associations with neuronal processes, including nervous system development and axon guidance (Fig. 6l). This suggests that AP-1 regulatory divergence contributes to species-specific neuronal programs.

Lastly, we asked which sequence changes drive this AP-1 response re-tuning across species. We identified the seqlets that contribute to the AP-1 response at human D1 peaks and asked which of them, where sequence has diverged across species, explained the most variation in the model’s predicted AP-1 response. Besides the general sequence divergence between species, the motif changes in total accounted for 13.8% of the predicted AP-1 response loss in non-human species. The AP-1 motif itself explained 12.2%, followed by a small set of TFs, including the activity-regulated cooperator EGR, MEF2^57^ (Fig. 6m). To understand why AP-1 responses are predicted to be less conserved than general chromatin accessibility, we also identified seqlets for ATAC signal predictions. Those seqlets have similar mutation rates as AP-1 across species (Extended Data Fig. 11e), but when mutated, they showed a smaller effect on chromatin accessibility prediction than the effect of AP-1 mutation on AP-1 response predictions (Extended Data Fig. 11f). The dominant contribution of the AP-1 motif makes the AP-1 response more vulnerable to evolution.

As an example, at a PLA2G4A/PTGS2-proximal D1 enhancer (Fig. 6n,o), the model’s predicted induction in humans, where the peak is strongly induced, is carried by four seqlets: two AP-1 sites, a JDP2 (bZIP) site, and an Nr2e1 site. In chimpanzee, marmoset, mouse, and elephant, single substitutions inside these motif cores, but not surrounding silent changes, explain the predicted loss of induction. A conserved AP-1 induction signal is read out into cell-type- and species-specific enhancer outputs through a small number of single-base changes at a defined set of cooperating TF sites. This cooperative motif logic is recoverable from sequence alone across an entire mammalian clade.

## Discussion

Cellular regulation is encoded in DNA but interpreted differently across cell states, developmental stages, activity programs, and species. Existing sequence-to-function models read regulatory information from DNA at high resolution, but most predict a fixed panel of tracks, one per cell type or tissue in the training data. Cell state is therefore something the model decides rather than something the user can specify. Single-cell foundation models take the opposite approach: they learn representations of cellular identity and state, but operate on measurable molecular features, and do not directly interpret DNA sequence or genetic perturbation. Bolero bridges these two frameworks by making the cell state, and the variables that modulate it, into inputs the user applies. An atlas-scale embedding, a developmental age, a species, and a continuous activity score each condition the network at every layer, so that chromatin accessibility and transcript abundance can be predicted from DNA in a specified cellular context. That conditioning is what makes Bolero substantially better than specialised models at telling cell states apart, and what allows a sequence to be evaluated in a state lying between the measured ones, or under a stimulus that was never applied to it. The prediction problem shifts from what a sequence does in one measured assay to how the same sequence is interpreted across a coordinate system of cell type, region, developmental stage, activity state, and evolutionary background.

Cell-state-specific regulation is better captured as a continuous geometry than as a catalogue of discrete cell-type labels. Bolero preserves the genomic resolution of the underlying sequence model while gaining across-state resolution. Its ability to resolve cortical regional gradients, human developmental trajectories, and closely related cellular states indicates that the cell-state embedding influences how sequence features are read. Unattainable experimental data are no longer an obstacle to regulatory element discovery and understanding: Bolero uses embeddings from unmeasured states to improve over nearest-training-cell baselines, so learned representations become extrapolable rather than purely memorised. This generalisation remains incomplete, especially for harder held-out settings where regulatory programs differ substantially from the training states. Still, it opens a practical route to prediction in sparsely sampled regions of the state space. No experimental atlas can exhaustively cover combinations of cell type, stimulus, age, genotype and species, so partial extrapolation across these axes is what makes predictive regulatory atlases achievable.

This cell-state resolution is particularly relevant for human genetics. Bolero was not designed solely as a variant-effect predictor, yet its QTL performance is comparable to current specialised models: variant effects can be predicted without giving up broad cell-state coverage. That coverage lets bulk QTL effects be projected onto plausible cellular origins, base-pair attributions be related to GWAS heritability, and non-coding de novo mutations be interpreted across developmental and lineage contexts. Non-coding variant interpretation then becomes a question of where and when a regulatory perturbation acts, not only whether it changes a regulatory element. The eQTL predictability analysis also marks where current sequence models fall short: promoter-proximal and active-chromatin variants are captured better than many distal variants, consistent with the difficulty of modelling long-range enhancer–promoter communication and broader trans-regulatory context.

Which enhancers respond to a shared activity signal is decided by their local cooperative transcription factor context. Neuronal activity induces a broadly shared AP-1-associated program, yet the enhancers that respond to this signal are highly cell-type-specific. A controlled stimulation experiment can be carried out in a model organism but not in a human, and yet cells in the body are heterogeneously stimulated all the time. The premise of the AP-1 score is that an unperturbed atlas already contains the result, spread across cells of the same type. Entering AP-1 activity as a conditioning variable shared across datasets asks the model to learn from natural heterogeneity rather than a stimulus someone applied, and Bolero-AP1-Score then separates the trans induction signal from the cellular context in which it is decoded. The model predicts activity-induced chromatin changes in behaviour-paired and sensory-experience paradigms held out from training, reproduces DG- and CA1-specific enhancer responses, and identifies local sequence features that tune AP-1 responsiveness. Directed sequence evolution further shows that the learned grammar is not only readable but writable. Across the human body, AP-1 responsiveness is organised by cell-class-specific cooperator vocabularies and by motif syntax, including spacing- and orientation-dependent interactions with cooperator motifs. These analyses convert AP-1 motif enrichment into a quantitative model of cooperation, in which motif identity, local spacing, and the cell-state-specific transcription-factor repertoire jointly determine the predicted response. Variation within a cell type, which a cell-type-level atlas averages away, becomes a set of cell-type-specific regulatory rules once it is given an axis of its own. Structural modelling provides hypothesis-generating support for possible DNA-mediated co-binding at some composite elements, but these predicted interactions remain to be tested experimentally.

Sequence conservation is a poor guide to whether a regulatory response is conserved. In basal ganglia neurons, AP-1-inducible responses are predicted to be retained across mammals less often than cell-type-defining accessibility, even though the two are written into elements of comparable sequence conservation. Reading that difference required predicting regulatory output directly from sequence, which extends comparative genomics from sequence conservation to predicted functional conservation. Matched single-cell regulatory atlases exist for only a small number of species, whereas genome assemblies now span a broad mammalian clade. Bolero can be queried on species genomes that lack matched functional data, predicting cell-type-specific accessibility and AP-1-associated responses across evolutionary distance. While it is not feasible (or ethical) to carry out the stimulation experiment in humans, Bolero nominates basal-ganglia human-biased enhancers at genes of nervous-system development and axon guidance, which makes activity-dependent regulation a candidate substrate for what is specific about the human brain. Our modelling indicates that species-specific tuning arises from single-base changes in AP-1 and cooperator motifs, but canonical motif-core disruption explains only a fraction of the within-peak, across-species variance in the AP-1 response. Inducible enhancer responses therefore appear to evolve quantitatively, through both recognisable motif changes and more distributed sequence features.

Despite the advances, we still have challenges and limitations. Many Bolero predictions were tested against independent experimental datasets, including activity-regulated, genetically labelled, and cross-species chromatin profiles, but the current readouts remain centred on accessibility and transcript abundance. These measurements capture regulatory consequences; they do not directly resolve transcription-factor occupancy, protein abundance, chromatin history, enhancer–promoter communication or three-dimensional genome organisation. Future models could extend Bolero-style cell-state conditioning to additional molecular modalities and perturbation-paired measurements, or carry the cell-state-aware approach onto more capable base models such as AlphaGenome. The AP-1 chromVAR score is likewise a proxy for one inducible transcription-factor axis; other signalling pathways, disease states, drug responses, genotypes, and environmental contexts will each need their own quantitative conditioning variables. At the model sizes and data budget we tested, scaling analyses show performance to be data-bound rather than compute-bound, so further gains should come from broader atlases, additional species, and richer matched measurements. Closing the experimental design loop matters too, both for testing predicted cis-regulatory grammar and for developing cell-state-targeted regulatory elements. The endpoint is a sequence-native predictive regulatory atlas that can be queried across cell state, perturbation, and species.

## Methods

### Model architecture

Bolero predicts single-cell-state-resolved chromatin accessibility directly from genomic DNA sequence. The model takes a 524,288 bp one-hot-encoded DNA sequence as input and produces 16,384 output bins at 32 bp resolution, with the flanking 16,352 bp cropped, following the Borzoi output geometry. The sequence backbone is the Flashzoi variant of Borzoi, comprising a convolutional DNA stem, a residual convolutional tower with a U-Net-style upsampling path, eight transformer layers (1,536 channels, 8 attention heads, FlashAttention), separable convolutions, and final joined convolutions feeding the output heads. The backbone weights are initialised from the published Flashzoi fold-0 checkpoint.

Cell-state conditioning is implemented as conditional low-rank weight generation^58^ added to the frozen backbone. For each adapted layer, the low-rank factors A and B are generated from the cell-state embedding by an embedding multilayer perceptron, so that the effective weight is W = W_base + (alpha / r) * A(e) @ B(e), where e is the per-cell-state embedding and the low-rank scale alpha is 0.02. B is zero-initialised, so the model begins as the unmodified backbone and departs from it only as conditioning is learned. Per-layer ranks r differ by module (convolutional DNA stem 2; residual tower and the first U-Net stage 6; transformer layers 30, with the attention query, key, value and output projections held non-conditional; horizontal and upsampling paths 30; separable convolutions 10; final joined convolutions 60; output head 1). The conditioning module concatenates a cell-state encoder, a condition encoder and a dataset-shared embedding. A dataset-shared embedding is a continuous representation or metric computed on one common scale across all datasets, so that its values are comparable between them rather than interpretable only within one; here it carries the AP-1 activity score. The only condition term used in the released model is Tissue (a low-dimensional condition embedding, for example, 2 dimensions for Tissue and 1 for Age). A non-conditional baseline configuration uses a single shared rank of 8 across all adapted layers.

The reference signal is a bulk accessibility profile, obtained by summing the single-cell data across cells and used unchanged for every cell state. It carries no cell-state-specific information, only a local, general signal for each region, which supplies scale information that DNA sequence alone does not. This reference signal can be injected into the DNA embedding at 32 bp resolution through a three-convolution signal encoder (zero-initialised) whose output is added to the sequence embedding. During training, the reference signal is withheld on 20% of training steps, which lets the same model run in a DNA-only mode at inference, with no reference signal provided.

The model exposes several output heads. The count (accessibility) head is a convolution followed by a softplus, trained with a Poisson-multinomial loss. A delta head predicts the paired difference in accessibility between a reference cell state and a target cell state: the change from one state to the other, not the absolute level in either. It is trained with a mean squared error loss, and predictions made through it are referred to below as delta mode. A hypernetwork head reproduces the published Scooby model, in which the cell-state embedding generates per-cell convolutional output weights, and is used as a baseline.

### Model training

Models were trained with AdamW^59^ (learning rate 5e-5, weight decay 1e-8, global gradient-norm clip 0.5) in bfloat16 mixed precision. The accessibility head used the Poisson-multinomial loss (total weight 0.16, epsilon 1e-7). The learning-rate schedule was a linear warmup over 500 steps followed by a polynomial decay (power 5). Each epoch used 5,000 training batches and 300 validation batches, batch size 4 with gradient accumulation of 2. No early stopping was used; the best checkpoint per epoch was retained.

Training followed a multi-stage curriculum: (1) an initialisation stage of 50 epochs; (2) a dataset-specific stage of 15 epochs in which only the conditioning encoders (the cell and condition embeddings) were trained; (3) a combination stage that merged the per-dataset conditioning parameters; (4) a final joint stage of 100 to 200 epochs; and (5) a gene-count head stage of 200 epochs. The per-cell-state embedding fed to conditioning was a 30-dimensional per-metacell latent, and the only condition term used was Tissue.

### Data collection and processing

The training atlas comprises 33 single-cell datasets totalling 10.7 million cells across 6 species: human (5.4M cells, 13 datasets), mouse (4.2M cells, 12 datasets), macaque (601K cells, 3 datasets), marmoset (424K cells, 3 datasets), bonobo (17K cells, 1 dataset) and opossum (21K cells, 1 dataset). Each dataset, its species, cell count, assay modality, cell-state embedding method and latent dimension, and source publication, is listed in Supplementary Table 1.

Peaks were called with MACS3^60^ through SnapATAC2^61^ and standardised to 500 bp. Metacells were computed with SEACells at a resolution of approximately one metacell per 75 cells, yielding roughly 159,000 metacells, with at most 3e6 fragments per metacell. Metacells overlap rather than partition the cells, so their number is larger than the cell count divided by that resolution. For training, pseudobulks were formed with a minimum of 1,500 cells and a minimum of 5e6 fragments per pseudobulk, with an oversampling factor of 2 and a maximum overlap of 0.1 between pseudobulks. For biological inference, pseudobulks were instead defined by biological groupings (subclass by age by region) with a target coverage of 5e6 or 25e6 fragments. Coverage tracks were computed at 32 bp resolution.

The genomic train, validation and test split followed the Borzoi fold-0 split (training folds 2 to 7, validation fold 0, test fold 1), corresponding to 695 held-out test windows of 524,288 bp each, about 11.7% of the genome.

Cell-state embeddings (30 latent dimensions) were produced with MultiVI for multimodal (RNA plus ATAC) datasets and PoissonVI for ATAC-only datasets. For the held-out generalisation analysis and the embedding benchmark, RNA-only scVI and scANVI embeddings (30 dimensions) and a STATE embedding (2,058 dimensions, zero-shot from the SE-600M model) were additionally computed.

### Prediction modes

The same trained Bolero was run in two inference modes, selected by a single flag rather than by retraining: a signal-aware mode supplies a reference accessibility signal (the reference cell state being a fixed set of 1,000 metacells identical across all pseudobulks), and a DNA-only mode withholds the reference signal entirely (the reference-signal dropout applied during training lets one model serve both). Per-peak accessibility was summarised as the sum of predicted signal over the peak (a 500 to 512 bp window). The count and hypernetwork heads were used without a signal channel; the delta mode predicts a target cell state against a supplied reference and scores the difference. Signal-aware absolute prediction was used for the accuracy benchmarks (Fig. 1, Extended Data Figs. 1 and 2, and Extended Data Fig. 3a,b) and state-specific reconstruction (Extended Data Fig. 2); the delta mode was used for the within-cell-type and held-out-generalisation analyses (Fig. 2, Extended Data Figs. 3d and 4); and DNA-only inference was used for the DNA-only benchmark comparison and for all cross-species prediction (Fig. 6, Extended Data Fig. 11). The model, training data, inference data and inference mode behind every main and Extended Data figure panel are listed in Supplementary Table 5, and the models, baselines and inference modes it names are defined in Supplementary Table 6.

### Pseudobulk accuracy metrics

Accuracy was computed on peaks by pseudobulk matrix. The across-peak (Profile) correlation is the per-pseudobulk Pearson correlation across peaks; the across-sample (Sample) correlation is the per-peak Pearson correlation across pseudobulks; the coefficient of determination R squared is reported per peak. To focus on informative regions, peaks were stratified into 5 quintiles by their standard deviation across pseudobulks, cumulatively, so that group 0 is all peaks and group 2 is the top 60% most variable peaks; unless noted, reported values are the group-2 mean plus or minus standard deviation. Benchmark evaluations used a target coverage of 5e6, 100 sampled pseudobulks, test regions only, and the fixed 1,000-metacell reference, and compared Bolero, a fine-tuned Borzoi baseline, the same Bolero run in DNA-only inference mode, and Scooby. Gene-expression prediction was benchmarked analogously on a genes-by-pseudobulks matrix (across-gene and across-sample Pearson correlation and R-squared), comparing Bolero and DNA-only Bolero with Decima. Per-dataset accessibility benchmark values for every model and metric are provided in Supplementary Table 2, and the corresponding gene-expression values in Supplementary Table 3.

### Differential accessibility regions

Differential accessibility regions (DARs) were called with pyDESeq2^62^ pairwise contrasts at adjusted p < 0.01 (Benjamini-Hochberg), with additional thresholds (minimum fraction of cells, absolute log2 fold-change) and the contrast and pseudoreplicate structure specified per analysis. DAR sets used for accuracy evaluation were intersected with the held-out test regions.

### Variant effect scoring

Variant effects were computed by substituting the reference and the alternate base into the one-hot sequence at the variant position, predicting each sequence, and taking the difference. For chromatin-accessibility QTLs (caQTLs), per-bin predictions were clamped at zero and summed over the bins of the variant’s peak, and the effect was the log2 ratio of alternate to reference peak sums. For expression QTLs (eQTLs), the RNA head was aggregated into a gene-level count over the gene body with a strand-aware gene mask, and the effect was the log2 ratio of alternate to reference gene counts. Conditioning used each dataset’s 30-dimensional MultiVI embedding; because each variant is scored as a within-cell-state reference-versus-alternate contrast, the embedding does not leak the effect. All scoring used the 524,288 bp window and was restricted to held-out test regions.

### Sequence attribution and motif annotation

Per-base contribution scores were computed with input-times-gradient (Captum), as the gradient of the summed peak prediction with respect to the one-hot input, stored over a 1,024 bp window centred on the site of interest. For the AP-1 score model, hypothetical contributions were defined as the difference between the AP-1-high and AP-1-low attributions. Seqlets were extracted with the tangermeme recursive-seqlet procedure (p < 0.001 for single-variant attribution; threshold 0.03 for the score-model motif-discovery seqlets; seqlet length 4 to 25 bp, 25 bp flank). Sequence motifs were annotated against the JASPAR database^63^, either by scanning candidate sequences with motifmatchr or, for de novo patterns, by matching with TomTom^64^.

### Accuracy across variability cutoffs, state-specific reconstruction and single-versus multi-dataset models

Prediction accuracy was examined as a function of feature variability on the held-out test fold of a representative accessibility dataset (Zhang 2021 whole-body atlas) and a representative expression dataset (human brain development), each evaluated at 5e6 target coverage with 100 pseudobulks sampled at random (the same fixed 1,000-metacell reference and signal-aware mode as the benchmark). Features (peaks or genes) were ranked by their standard deviation across the 100 pseudobulks and binned into quintiles; cumulative cutoffs retained the top 100%, 80%, 60%, 40% and 20% most variable features, and at each cutoff the across-peak/across-gene correlation and R squared (one value per pseudobulk) and the across-sample correlation and R squared (one value per feature) were recomputed, so that accuracy could be read as a function of how variable the retained features are.

To visualize state specificity directly, features were first restricted to the top 40% most variable, and the most state-specific among them were selected by taking, for each pseudobulk, the 500 features with the highest fold-change of that pseudobulk over the mean of all others, then pooling and de-duplicating across pseudobulks (24,768 peaks for the accessibility dataset, 456 genes for the expression dataset); measured and predicted values were log1p-transformed and z-scored per feature and shown as paired heatmaps with pseudobulks ordered by hierarchical clustering of the measured data.

To weigh the benefit of joint training, each dataset’s accuracy under its own single-dataset low-rank model was compared against the accuracy on that dataset of the multi-dataset Bolero model trained on all datasets of the same species (WholeHuman or WholeMouse), over the top 60% most variable features, across the four metrics (one point per dataset subset; 27 subsets from 18 human and mouse datasets).

### Model scaling, conditioning ablation and embedding selection

To assess capacity, the conditional model was retrained at six sizes by setting the hidden dimension to 64 times a scaling factor of 1 to 6, giving trainable-parameter counts from about 0.1e9 to 3.5e9 (baseline 1.01e9), each on a fixed 600,000-sample subset, and accuracy was tracked over training and across peak-variability groups; it plateaued near 1e9 parameters. At each training checkpoint and model size, variant effects were also scored on the adult-neuron caQTL set (n = 673) and summarised as the signed and unsigned correlation with the measured effects. To localise where conditioning acts, a per-layer ablation made conditioning active in only one module at a time, from the convolutional DNA stem through the output head, and compared each against the fully conditional model. To select the cell-state embedding, scVI, scANVI, STATE (zero-shot), PoissonVI and MultiVI were compared on the HumanBrainDev developmental dataset with a delta prediction task, restricted to peaks with a standard deviation above 5 across pseudobulks, for Bolero, DNA-only Bolero and Scooby (Extended Data Fig. 3d).

### Regional and developmental accessibility

To test whether Bolero captures accessibility differences between closely related cell states, predictions were made in delta mode with the reference signal supplied, using the count-loss checkpoint as Bolero and Scooby as the baseline, at 5 Mb coverage and keeping pseudobulk groups with more than 5e5 fragments. Pseudobulks were defined as subclasses by brain region in adult mouse cortex and as subclasses by developmental stage in human brain development. Accuracy was evaluated specifically on differential peaks, as the Pearson correlation between measured and predicted accessibility was computed for each peak across these pseudobulks. In adult mouse cortex, DARs were called between subclasses within each brain region and between brain regions within each subclass (minimum fraction of cells 0.05, absolute log2 fold-change at least 0.25), giving 36,411 subclass DARs and 22,871 brain-region DARs after intersection with test regions (from full unions of 301,895 and 187,119). In human brain development, DARs were called with donor pseudoreplicates between subclasses within a developmental stage and between developmental stages within a lineage, giving 43,399 subclass DARs and 20,030 developmental-stage DARs after intersection with test regions.

### Held-out cell-state generalisation

A central question is whether Bolero predicts the accessibility of cell states it never saw in training, rather than memorising trained states. To test this directly, cell states were systematically withheld along the biologically meaningful axes of the human developmental atlas: by developmental time (holding out a trimester), by cell-type subclass (holding out a subclass), by lineage age (holding out an age within an inhibitory lineage) and by cortical region (holding out region-specific subclasses), as well as a random holdout of cell states. For each of 24 held-out cases (eight held-out configurations, each at three held-out fractions or subgroups a/b/c), a model was trained only on the retained cell states and then asked to predict the held-out states from sequence. Its accuracy was compared with a Closest baseline that simply copies the prediction of the nearest training cell state (the nearest training pseudobulk by Euclidean distance in the embedding space, reusing its embedding and condition). A model that truly generalises from sequence should beat this nearest-neighbour copy.

Because the cell-state embedding is derived from the same cells’ multiome data, a concern is that ATAC information in the embedding, rather than the sequence, drives the prediction. To exclude this, the entire analysis was repeated with RNA-only embeddings (scVI and scANVI), alongside the multimodal MultiVI embedding and a STATE embedding. Accuracy was measured as the mean over peaks of the per-peak across-sample Pearson correlation, on a panel-specific set of differential peaks and cell states with coverage above 5e5 fragments, with a per-case all-data model as a positive-control ceiling. The main Fig. 2g–j panels feature 18 of the 24 cases across four representative axes with panel-shared differential-peak sets and the scVI embedding, over which Bolero beat its own Closest baseline in 17 of 18 (the one exception a held-out-timepoint case). Over all 24 cases with per-case differential-peak sets (Extended Data Fig. 4), Bolero beat Closest in 22 of 24 (scVI), 21 of 24 (scANVI) and 23 of 24 (MultiVI); because the two analyses use different pseudobulk and peak selections, the same case can rank differently between them. Performance was essentially unchanged across embeddings (RNA-only within about 0.04 of MultiVI), showing the result does not depend on ATAC information in the conditioning embedding. The STATE embedding was weak and had no native Closest baseline.

### caQTL and eQTL prediction

Variant effects were scored as in the shared section and compared with measured quantitative-trait-locus effects. Adult-neuron caQTLs (Zeng 2024, fine-mapped by SuSiE^65^ at posterior inclusion probability above 0.5; n = 673) were scored, and effect prediction was evaluated both as the signed correlation with measured effects (Pearson r = 0.79) and as the AUROC for prioritising fine-mapped caQTLs (673) against background variants (1,094), comparing Bolero (all-data and brain-only), Scooby and ChromBPNet (ENCSR868FGK bias model). Whole-blood eQTLs (GTEx^66^, posterior inclusion probability above 0.9; n = 851) were scored on RNA pseudobulks with a signed correlation of Pearson r = 0.38, and blood-eQTL prioritization was benchmarked across 19 immune cell types (OneK1K, posterior inclusion probability above 0.9; 957 positive and 21,617 negative variants) as the average precision per cell type, comparing Bolero (all-data and blood-only), Decima, Borzoi and TSS distance. Tissue specificity was examined by correlating Bolero’s predicted eQTL effects in each ENCODE Human Atlas cell type with the observed effects of each GTEx tissue’s eQTLs.

### Cell-type deconvolution of bulk eQTLs

To resolve the cell types underlying a bulk-tissue eQTL, each fine-mapped whole-blood GTEx eQTL was scored separately in every cell state of a peripheral-blood mononuclear-cell (PBMC) reference, taking the effect as the log2 ratio of predicted gene counts between the alternate and reference alleles. This yielded, for each eQTL, a vector of predicted effects across blood cell types, and a bulk-level prediction was recovered by averaging these per-cell-type effects weighted by each cell type’s abundance in the reference. The eQTLs with an appreciable predicted effect (absolute abundance-weighted log2 fold-change above 0.01; 437 of 851) were displayed as an eleven-cell-type-by-eQTL heatmap with eQTLs hierarchically clustered (Extended Data Fig. 6a). The deconvolution was validated at cell-type resolution against an independent single-cell eQTL cohort: GTEx eQTLs were intersected with eQTLs fine-mapped to individual cell types in OneK1K (posterior inclusion probability above 0.9), and for each overlapping association (73 gene-variant pairs, expanded to 178 cell-type-specific associations after mapping OneK1K cell types onto the eleven reference cell types) Bolero’s deconvolved effect in the matched cell type was compared with the measured cell-type-specific effect size, agreeing in sign and magnitude (Pearson r = 0.44, Spearman ρ = 0.52; Extended Data Fig. 6b).

### eQTL predictability

To characterise which eQTLs the model captures, eQTLs with a measured effect of at least 0.1 in absolute value were labelled predictable when the cell-abundance-weighted predicted log2 fold-change exceeded 0.1 in absolute value with the same sign (180 eQTLs), and unpredictable when the absolute predicted log2 fold-change was at most 0.1 (651 eQTLs); the remaining 20 eQTLs (opposite-sign predictions or a measured effect below 0.1) were not assigned. Predictable variants were tested for enrichment in genomic categories (Annovar RefGene^67^, Fisher exact with Benjamini-Hochberg correction), for active-histone-mark signal (FAVOR annotations^68^, Mann-Whitney U), for distance to the TSS (predictable median 176 bp versus 2,113 bp, Mann-Whitney p = 3.3e-19) and for SpliceAI score (mean 0.0532 versus 0.0302, Mann-Whitney p = 7.0e-3). The same analysis reports the adult-glia caQTL AUROC and a single-dataset versus whole-human versus 10M model comparison (Extended Data Fig. 5a–c).

### Attribution-based GWAS heritability enrichment

To ask whether the sequence features Bolero relies on are heritability-relevant, per-base attributions were computed for the merged developmental neuronal class and used to partition accessible SNPs into a high-attribution set (peak positions overlapping seqlets) and a low-attribution set (peak positions outside seqlets). The high-attribution set comprised 0.5%, and the low-attribution set 3.6% of all SNPs tested. The two annotations were tested jointly against baselineLD by stratified LD-score regression^29,30^ (S-LDSC; baselineLD v2.2, 1000 Genomes EUR LD^69^, hg38 lifted to hg19, overlap-annot), and per-trait coefficient z-scores were converted to p-values and Benjamini-Hochberg corrected across traits; 50 traits passed coefficient FDR < 0.05 (two-sided), with brain and cognition traits (schizophrenia, bipolar disorder, ASD, ADHD) highlighted.

### De novo variants in autism

To test cell-type-resolved interpretation of disease variants, de novo variants from the Zhou 2019 autism cohort (lifted hg19 to hg38; 127,137 SNVs; 65,146 from probands and 61,991 from siblings) were scored across 149 developmental pseudobulks. A variant was called large-effect when its absolute log2 fold-change exceeded the per-pseudobulk 99.9th percentile, and proband-versus-sibling enrichment of large-effect variants was tested per cell class and developmental stage by one-sided Fisher exact tests with Benjamini-Hochberg correction. The same test was stratified by gene category, comparing all genes (n about 126,039 SNVs) with SFARI autism genes (n about 30,386) and loss-of-function-intolerant genes (ExAC pLI > 0.9; n about 53,067), using gene windows from 250 kb upstream of the TSS to 50 kb past the TES, with a Wilcoxon signed-rank test across pseudobulks. As a baseline without selection on predicted effect (labelled Random in Fig. 3h), the case-to-control odds of de novo variants falling within each gene category was computed with a one-sided Fisher exact test; this baseline takes one value per gene category. Individual disease-associated variants, such as one near ZBTB20 that introduces a SOX core motif, were dissected with per-base attribution and with predicted accessibility and gene-expression changes resolved per metacell across the developing-brain atlas.

### Single-variant attribution across the atlas

To show that a single variant’s effect can be resolved across many cell states at once, two example variants were scored across the full atlas of pseudobulks. For each, the reference-minus-alternate per-base attribution over a 200 bp window was computed in every pseudobulk and shown as a heatmap of the per-position attribution change ordered by hierarchical clustering (Ward) of the pseudobulks, alongside the reference, alternate and difference per-base attribution logos, resolving the cell states in which the variant reshapes local sequence attribution.

### AP-1 activity-score model

To predict stimulus-dependent accessibility, an activity-score model was built in which an AP-1^36^ activity score (the z-scaled chromVAR deviation for the JASPAR motif MA0478.2) is supplied as an additional per-pseudobulk conditioning input alongside the cell-state embedding, concatenated into the conditioning module through a shared encoder (layer normalisation). The score model was fine-tuned from the pretrained 10M Bolero checkpoint with the same low-rank trainer and count head for 200 epochs across the full atlas. The atlas carries the control cells of the activity datasets only: their stimulated and perturbed cells were excluded from both pretraining and this fine-tuning, so the model learns the cell-state embedding space of those datasets without ever seeing activity-induced chromatin, and the activity paradigms used for evaluation are held out. Predictions were validated against measured activity-induced differential accessibility across five paradigms (fear engram, stress, novel environment, mouse visual experience and human visual experience), quantified against a motif-scan baseline by an odds ratio (agreement between the predicted and measured up-versus-down direction among the significant differential peaks) and an average precision (recovery of the significantly up-regulated peaks against all other peaks). Predictions were made in DNA-only mode for one control or early pseudobulk of the evaluated cell type, at a fixed grid of AP-1 conditioning values (-3, -1.5, 0, 1, 3, 5 and 8 in z-scaled chromVAR units), and the predicted effect was the log2 ratio of the signal at a high AP-1 value to that at a low AP-1 value. The measured contrast and the two AP-1 values differed by paradigm. For the fear engram, DG Fos-Trap+ cells were compared with Trap− cells of the 24-h group, and the prediction for the 24-h Trap− pseudobulk compared AP-1 values of -1.5 and +8. For novel-environment exposure, the highest and lowest AP-1-score groups of CA1 cells within the 1-h exposure sample were compared, and the prediction for the control CA1 pseudobulk compared - 1.5 and +5. For stress, the highest and lowest AP-1-score groups of CA1 cells within the 15-min restraint sample were compared, and the prediction for the 45-min handling CA1 pseudobulk compared -3 and +1. For mouse visual experience, L2/3 IT neurons of the (33.5, 35.5] age group were compared with those of the (25.5, 31.5] age group, and the prediction for the earlier group compared -1.5 and +5. For human visual experience, postnatal L2/3 IT neurons were compared with prenatal ones, and the prediction for the prenatal pseudobulk compared -1.5 and +5. Measured differential peaks were the DESeq2^62,70^ contrasts with an absolute log2 fold-change above 0.5, except for stress, where no fold-change cutoff was applied. Evaluation was restricted to peaks above the 40th percentile of total measured accessibility (the 60th for stress). Predictions were quantile-normalized across AP-1 values for the three hippocampal paradigms but not for the two visual-cortex paradigms. Peaks were annotated for an AP-1 motif hit by a motifmatchr scan of MA0476.2 (FOS; JASPAR2024, p = 5e-5).

The choice of AP-1 as the activity score was justified directly from the measured data. For each paradigm, the per-cell chromVAR deviation z-scores of all JASPAR motifs were computed, and the change between the stimulated and control conditions was tested per motif by a Mann-Whitney U test (Benjamini-Hochberg corrected, requiring at least 10 cells per group); the AP-1 motif cluster showed the largest deviation change in every paradigm. The AP-1 chromVAR deviation also increased monotonically along the visual-cortex developmental trajectory in both mouse and human. It separated activity-labelled (Trap+) from unlabelled (Trap−) neurons in a cell-type-specific manner across hippocampal subclasses and timepoints after fear learning (Extended Data Fig. 7). To test cell-type specificity, the model was conditioned either on an embedding computed from only the non-stimulated control cells or on a broad brain-atlas embedding, so that the AP-1 score alone carried the stimulus signal, and predictions were examined on dentate-gyrus-specific, shared and CA1-specific peaks and at individual loci, with the underlying motifs identified by attribution.

### Enhancer design by directed evolution

Synthetic enhancers were designed to realise target cell-type-specific accessibility patterns (for example, introducing an AP-1 response in dentate gyrus by adding bHLH, NF-I and AP-1 motifs, or suppressing accessibility in CA1 by removing bHLH and AP-1 motifs). The design used beam-search random mutagenesis with selection: starting from a reference sequence, each round produced candidate sequences by applying random point mutations within the evolution window to the current parents (number of mutations equal to the mutation rate times the window length, mutation rate 0.0034, which gives one mutation per round, each mutation forced to change the base), then reverting a fraction of the introduced differences toward the reference (back-mutation probability 0.1, applied only when at least 3 positions differed) to keep designs close to the start. The current parent was included in every batch so a non-improving round could be discarded; duplicates were filtered with a cache; the model scored all candidates against the target objective, and the top-scoring sequences became the next round’s parents. Each design target was pursued in 50 independent runs of 15 rounds, scoring 64 candidates per round (about 48,000 sequences per target); a worked example peak was evolved toward six target accessibility states (defined as the predicted accessibility in CA1 and dentate gyrus at AP-1-low and AP-1-high). The final designs carried roughly 0.33% to 2% mutated positions.

### De novo motif discovery

To learn the sequence grammar of AP-1-dependent accessibility, attribution peaks were selected per pseudobulk (above the 99.5th log-fold-change percentile and the 80th mean percentile, at most 15 regions per pseudobulk across 1,204 pseudobulks) and their hypothetical contributions passed to TF-MoDISco^43^ (modiscolite v2.4.0; up to 200,000 positive seqlets, 400 bp output window, sign tracking), yielding 86 positive and 1 negative patterns curated to 22 used patterns. Patterns were matched to known motifs with TomTom against JASPAR2024 (p < 0.01; the motif-to-transcription-factor-family cluster mapping used for annotation is given in Supplementary Table 7), and instances were called genome-wide with finemo-gpu v0.30 (CWM trim 0.4, global lambda 0.5, minimum hit correlation 0.5, up to 5 rounds; hits retained at correlation > 0.8). To visualise the motif repertoire, the representative seqlets were embedded in two dimensions by t-SNE (openTSNE) computed on a modiscolite seqlet-affinity that compares seqlets by their per-base contribution scores, combining a coarse cosine similarity over nearest neighbours with a fine continuous-Jaccard alignment across sliding offsets and both strands; points were coloured by their de novo cluster, and the same embedding was recomputed separately for each example cell type’s own seqlets to display per-cell-type repertoires (so the coordinates differ between cell types).

### AP-1 motif cooperativity

To find motifs that cooperate with AP-1, AP-1-response peaks were defined per pseudobulk from the predicted AP-1-high-versus-low differential (response above the 95th percentile with the AP-1-high prediction above the 80th percentile; non-response below the 5th percentile), and each of the 22 de-novo motifs was tested for enrichment in response versus non-response peaks (two-sided Fisher exact with Benjamini-Hochberg correction; 164 pseudobulks by 22 motifs, about 932 significant at q < 0.05 and odds ratio > 1), with the AP-1 family dominant. To ask how broadly the AP-1 response programme is shared, genome-wide predicted AP-1 log fold-change vectors were correlated (Pearson) across the 164 human-body pseudobulks of the Zhang 2021 whole-body atlas on 100,000 sampled abundant peaks and Ward-clustered into cell-class groups. To measure cooperativity directly, motifs were inserted into shuffled backgrounds, and the predicted AP-1 response was read out as a function of spacing. The backgrounds were 50 accessible peaks selected for moderate accessibility and low cross-cell-type variability (above the median predicted accessibility and below the 10th percentile of absolute cross-cell-type log fold-change); each peak’s reference sequence was dinucleotide-shuffled (a single shuffle reused across all spacings and orientations of that peak), and any endogenous AP-1 site that arose by chance was removed by iteratively shuffling every occurrence of the canonical TGA-N-TCA motif and its single-mismatch variants. Into each background, the AP-1 consensus (TGACTCA) was inserted alone and paired with each cooperator consensus, across the four AP-1/cooperator strand orientations and the spacing grid (a sparse grid out to 150 bp, and a dense single-base grid of 1 to 30 bp to resolve the helical-phase periodicity; the figures show the dense grid). For each inserted construct, the effect was the log2 fold-change between the model’s predictions at the AP-1-high and AP-1-low activity-score conditions (score +6 versus -1.5; epsilon 1e-5) after per-pseudobulk quantile normalisation, summarised as the mean over the 50 backgrounds and two AP-1-head orientations for each spacing, orientation and pseudobulk, and tested against zero by one-sample t-tests with Benjamini-Hochberg correction. To relate cooperativity to cooperator availability, the FOX–AP-1 effect at the −4 bp spacing was summarised per cell subclass as the one-sample t-statistic against zero. RNA expression of FOX-family transcription factors was taken from a curated compendium of human cell-type expression^71^, and correlated (Pearson) with these t-statistics across the 41 cell subclasses with matched RNA; the five most positively and five most negatively correlated FOX genes are reported. The spacing-dependent response was reported across human-atlas cell types (Fig. 5) and broken down across the seven major cell classes of the human whole-body atlas (Extended Data Fig. 9).

### Structural modelling of TF cooperativity

To seek structural support for the predicted cooperativity, AP-1 dimers (FOS, FOSB, FOSL1, FOSL2 paired with JUN, JUNB, JUND) were folded against 218 cooperator proteins with AlphaFold3 (12,666 designs, including ternary protein-DNA-protein complexes carrying the AP-1 and cooperator motifs at a range of edge-to-edge spacings and binary protein-only, no-DNA controls). Candidate interactions were ranked by domain-level confidence (ipTM, pTM, predicted aligned error, contact probability and pLDDT). For the AP-1-TEAD/YAP1/TAZ (Hippo) axis featured in Extended Data Fig. 10, the inter-chain predicted aligned error between the AP-1 subunits (FOS, JUN) and the partner was read out along the AP-1 amino-acid sequence, contrasting predictions with and without DNA and as a function of the AP-1-TEAD motif spacing, with the DNA-binding domains annotated.

### Cross-species accessibility and AP-1 response prediction

To test sequence-driven prediction across species, the 10M Bolero count head was run in DNA-only mode (cell identity entering only through the metacell embedding and conditioning), summarising each 500 bp peak to a single value, on hg38, mm10, rheMac10, panTro6, calJac4/mCalJac1, monDom5 and panPan3. Cross-species correspondence was established by UCSC liftOver (minMatch 0.1 for pairwise cortex and basal-ganglia comparisons; minMatch 0.5 over TOGA^72^ all-filled chains for the cross-mammalian analysis), with orthologous peaks resolved by largest overlap. For the motor-cortex reconstruction across 20 shared subclasses, models trained on all mouse pseudobulks (WholeMouse) or all human pseudobulks (WholeHuman) were each applied to the human peak sequences and compared, per DAR peak, against measured human accessibility, measured mouse accessibility at lifted coordinates and the 10M model, testing whether weights trained in one species transfer to another from sequence alone. Accuracy was assessed on differential peaks and stratified by a conservation grouping that combined liftover success with terciles of the measured cross-species correlation.

To predict AP-1-dependent accessibility, each sequence was scored twice with the AP-1 conditioning value clamped low (-1.5) and high (+6) in z-scaled chromVAR units, and the AP-1 log fold-change taken as log2((s_high + 0.01) / (s_low + 0.01)), in the D1 medium spiny neuron population of each species. Ground-truth AP-1 differential accessibility was defined by splitting cells into terciles of AP-1 chromVAR deviation within each major class and sample and calling DARs with pyDESeq2 (high versus low tercile, minimum fraction of cells 0.05, more than 50 cells per group; AP-1-responsive at adjusted p < 0.05, absolute log2 fold-change > 0.5, above the 40th-percentile base mean). Per-species AP-1 prediction was evaluated against each species’ own DARs by average precision and odds ratio, against a motif-scan baseline (motifmatchr scan of MA0478.2, p = 5e-5) and a human-only model, in macaque (n = 120,416 peaks, 11,662 AP-1-up) and mouse (n = 76,129, 7,032 AP-1-up).

To test conservation across mammals, predicted D1 accessibility (the 10M Bolero model without AP-1 input) and the predicted AP-1 response (the AP-1 score model) were computed at every human peak (1,354,545 hg38 peaks) in 83 non-human mammals selected from the assemblies with TOGA alignment chains (the ten highest-quality assemblies per clade by contig and scaffold N50). Fig. 6j shows human and the 74 of these species into which at least 500,000 human peaks were lifted. Each peak was carried to a species by liftOver (minMatch 0.5) and predicted from that species’ own genome, so that the only information taken from human was the location of the orthologous region. Sequence conservation was quantified from the Zoonomia 241-way mammalian alignment as the fraction of aligned bases per peak; this alignment-fraction measure, not phyloP^73^ or phastCons^74^, is the conservation control for the retention comparisons below. Separately, per-peak mean Zoonomia phyloP (241-way mammalian) was compared between AP-1-responsive D1 peaks (n = 15,120) and other D1-accessible peaks (n = 208,718) by a two-sided Mann-Whitney U test (Extended Data Fig. 11d).

Functional conservation was scored against calls fitted on the human prediction. The accessibility call was the Youden-optimal threshold separating the 301,975 peaks detected in D1 medium spiny neurons (majorclass STR_D1_MSN; at least 0.05 fragments per cell in the low or high AP-1 tercile) from the 1,050,671 other peaks from the HMBABasalGangliaHuman datasets (AUROC 0.911; threshold 2.615; sensitivity 0.821; specificity 0.853). The response call was the Youden-optimal threshold on the predicted AP-1 log2 fold-change in human separating the 38,302 AP-1-up D1 peaks from the 263,673 other D1 peaks (AUROC 0.857; threshold 0.329; sensitivity 0.754; specificity 0.833). The retention of a peak was the fraction of the species into which it lifted whose prediction exceeded the threshold, among peaks lifted to at least 10 species. Retention of AP-1-inducible versus cell-type-defining peaks was compared across the 13 major cell types with a well-separated AP-1 prediction (at least 100 AP-1-up peaks, an AP-1-up-versus-background AUROC of at least 0.80 and a non-negative Youden threshold in the human prediction); AP-1-inducible peaks were retained less across species in all 13.

### Motifs and gene ontology associated with cross-species variation in AP-1 response

The motif and Gene Ontology analyses used 10,385 AP-1-responsive peaks selected on measured data: every peak of the human D1 differential-accessibility contrast between the lowest and the highest AP-1 score group with an absolute log2 fold-change above 1.5 and an adjusted p below 0.05. For Fig. 6l, these peaks were divided into three groups by k-means clustering (k = 3) of their predicted AP-1 log2 fold-change across species and the missing values were filled with the mean of the peak. The groups were ordered by their mean predicted response and termed conserved (1,178 peaks), primate-specific (3,004) and human-specific (6,203). Peaks were assigned to genes by GREAT single-nearest-gene rule^75^ (each peak assigned to the gene with the nearest transcription start site within 1 Mb) and tested for Gene Ontology biological-process enrichment against the genes detected in at least 10% of D1 metacells, with Benjamini-Hochberg correction applied within each peak set.

To identify the motifs behind the loss of the AP-1 response (Fig. 6m and Extended Data Fig. 11e,f), per-base attribution of the 10,385 peaks was computed with the AP-1 score model at the AP-1-high value in the STR-D1-MSN CB pseudobulk over a 1,024 bp window (10,375 windows lie inside a chromosome). Seqlets were called from the attribution and kept at an absolute seqlet attribution of at least 5, and each seqlet was annotated with its best-matching JASPAR matrix and the motif cluster of that matrix (879 matrices in 306 clusters). For the comparison with chromatin accessibility, a second peak set was needed, because at AP-1-responsive peaks the accessibility predicted without AP-1 input itself still tracks the AP-1-high prediction of the score model (median per-peak r across species 0.711, against 0.495 for the AP-1-low prediction, over 6,917 peaks). The accessibility set comprised 7,724 D1 peaks that are neither AP-1-up nor AP-1-down (adjusted p < 0.05 and absolute log2 fold-change > 0.5). They were drawn without replacement from the 238,579 such peaks so that each cell of a 20 × 20 grid on the joint quantiles of measured strength (log2 base mean) and predicted strength (log2 human prediction) held as many peaks as it held AP-1-responsive peaks (standardised mean differences −0.076 and −0.017). These peaks were attributed using the model without AP-1 input by the same procedure (17,547 seqlets with an absolute attribution of at least 5).

Motif change in each species was scored from the TOGA human-referenced chains, which give the aligned species base for each human base. Position weight matrices were converted to frequencies with a pseudocount of 0.1 distributed in proportion to the background base composition, which places every matrix on one log2-odds scale. For each seqlet, its matched matrix was placed on the human sequence, with position and strand chosen on human before the species was examined. The species window was centred between the nearest aligned human bases on either side of the motif and read as contiguous species sequence. The human and the species sequence were then scanned over the same shifts, up to one motif length on each side, so that the gain from searching cancels in the difference, and motif change was the human score minus the species score. A copy with no aligned base in its window was called lost (16.4% of AP-1 copy × species rows). Each peak × species pair was summarised, per motif cluster, by three terms: the change at the worst surviving motif copy, the mean change over surviving motif copies and the number of lost motif copies. When every copy was lost, the two change terms were set to 0, and the loss count carried the information. Substitutions per base at motif sites (Extended Data Fig. 11e) were counted over the seqlet span in the same alignment, over the 81 species that have both model predictions, a motif score, and the distance covariate discussed below; a motif found on both peak sets was pooled over its sites on both. Over all attributed motif sites, the rate was 0.156 at the AP-1-responsive peaks and 0.148 at the accessibility peaks, and single clusters ranged from 0.109 (CTCF) to 0.190 (EGR), with AP-1 at 0.152.

The loss of a call owing to motif change was estimated at each peak called positive in humans with a prediction in at least 20 species. For the response, these were the 8,780 attribution peaks with a predicted AP-1 response above 0.329 in humans; 8,455 of them carry an AP-1 seqlet, and 8,769 carry at least one attributed motif. For accessibility, these were the 6,657 matched peaks with a predicted accessibility above 2.615, of which 5,321 carry at least one attributed motif. Because both the motif terms and the loss of a call grow with evolutionary distance, each peak was given its own slope on a background divergence of each species, read from sequence that carries no attributed motif: B(s), the mean over peaks of the share of flanking bases (the 524 bases of the 1,024 bp window outside the 500 bp peak) that are not identical to human in the chain alignment. At each peak p, the call y_p_(s) (1 kept, 0 lost) was fitted across the species of the peak by a linear probability model, y_p_(s) = b_p_ + w_bkg,p_ B(s) + w_motif,p_^T^ x_p_(s), where x_p_(s) are the three motif terms, b_p_ is the intercept, and w_bkg,p_ and w_motif,p_ are the slopes; all coefficients were fitted separately at each peak. A motif term that took the same value in all species of a peak was omitted from that peak’s fit. The motif terms are 0 for a motif identical to human, so the fitted probability of keeping the call with an intact motif is b_p_ + w_bkg,p_ B(s), and w_motif,p_^T^ x_p_(s) is the reduction in this probability that the fit assigns to the motif change in species s. The motif loss of a peak is the mean of this reduction over the n_p_ species of the peak, L_p_ = (1/n_p_) Σ^s^ [−w_motif,p_^T^ x_p_(s)], which estimates the fraction of species in which the motif change removes the call. L_p_ was set to 0 where the call was the same in every species or where no motif term varied across species; a motif change shared by every species of a peak is therefore not attributed to the motif. Each motif cluster carried by at least 100 peaks was fitted alone at the peaks that carry it (21 clusters for the response and 29 for accessibility).

Two averages of L_p_ were reported.

1. The effect of a motif change where it occurs (Extended Data Fig. 11f) is the mean over the set P_k_ of peaks that carry cluster k, E_k_ = (1/|P_k_|) Σ_{p∈Pk}_ L_p_. AP-1 change removed the response call in 12.6% of species at the 8,455 peaks that carry AP-1, whereas the largest value for the accessibility call was 4.7% (EGR, 558 peaks).
2. The contribution of a motif to the conservation of the whole peak set (Fig. 6m) divides the same sum by the number of positive peaks in the set, C_k_ = (1/|U|) Σ_{p∈Pk}_ L_p_, so that it combines how often the motif occurs with its effect where it occurs.

Because AP-1 is carried by 8,455 of the 8,780 responsive peaks, its contribution, 12.2% (95% interval 11.8 to 12.6), is close to its effect where present. The contribution of all motifs together was computed in the same way from a joint fit at each peak, in which every motif cluster carried by the peak had its own three terms, and was divided by the same number of positive peaks. Over the 8,780 AP-1-responsive peaks, in which the response call was lost in 61.2% of species ((1/|U|) Σ_{p∈U}_ (1 − ȳ_p_), where ȳ_p_ is the fraction of species that keep the call), all motifs together removed the call in 13.8% of species (13.3 to 14.2). Over the 6,657 accessibility peaks, in which the accessibility call was lost in 26.2% of species, all motifs together removed the call in 3.5% (3.0 to 3.9). Permuting the motif terms between the species of each peak gave values between −0.3% and 0.1% for these totals. Intervals are 95% percentile intervals from 1,000 bootstrap resamples of peaks. For a contribution, peaks were resampled from all positive peaks of the set, with L_p_ = 0 for peaks without the motif; for an effect where the motif occurs, they were resampled from the peaks that carry the motif. A substitution rate is the total number of substituted bases divided by the total number of motif bases, so both totals were recomputed in each resample before the ratio was taken. Species were not resampled, so the intervals reflect variation between peaks.

## Data Availability

The Bolero-10M atlas, its pseudobulk count matrices and peak sets, including the datasets generated for this study, are available at https://huggingface.co/datasets/arcinstitute/bolero-data and are distributed with the bolerodata package (https://liuhlab.github.io/bolerodata/). The 33 source datasets, with their species, cell counts, source publications and repositories, are listed in Supplementary Table 1. This study also used data from GTEx (https://gtexportal.org/), OneK1K (https://onek1k.org/), ENCODE (https://www.encodeproject.org/), the 1000 Genomes Project (https://www.internationalgenome.org/), Zoonomia (https://zoonomiaproject.org/), TOGA (https://genome.senckenberg.de/download/TOGA/), JASPAR2024 (https://jaspar.elixir.no/), SFARI Gene (https://gene.sfari.org/), ExAC (https://exac.broadinstitute.org/) and FAVOR (https://favor.genohub.org/).

## Code Availability

Bolero is available at https://github.com/liuhlab/bolero and documented at https://liuhlab.github.io/bolero/; the bolerodata package is at https://github.com/liuhlab/bolerodata and documented at https://liuhlab.github.io/bolerodata/. Trained model weights are available at https://huggingface.co/arcinstitute/bolero-models.

## Acknowledgements

H.L. was supported by the Harvard Society of Fellows. M.E.G. was supported by funding from NINDS R01 NS115965 and Harvard Data Science Initiative Competitive Research Fund. S.M., T.G., Z.Y., J.Z. are supported by funding from the Arc Institute. We thank members of the Greenberg laboratory and Zhou laboratory for helpful discussions on the manuscript.

## Author Contributions

H.L., J.Z. and M.E.G. conceived the study. H.L., J.Z., R.Z. and J.D.B. developed the model. H.L., S.M., T.G., Y.C. and Z.Y. assembled the data. I.V. and L.T. generated the new single-cell datasets for the activity paradigms. H.L., S.M., T.G., Y.C., Z.Y. and J.Z. performed the analyses. H.L., S.M., J.Z. and M.E.G. wrote the manuscript, and all authors revised it. H.L., J.Z. and M.E.G. supervised the study.

---

**Extended Data Figure 1.**
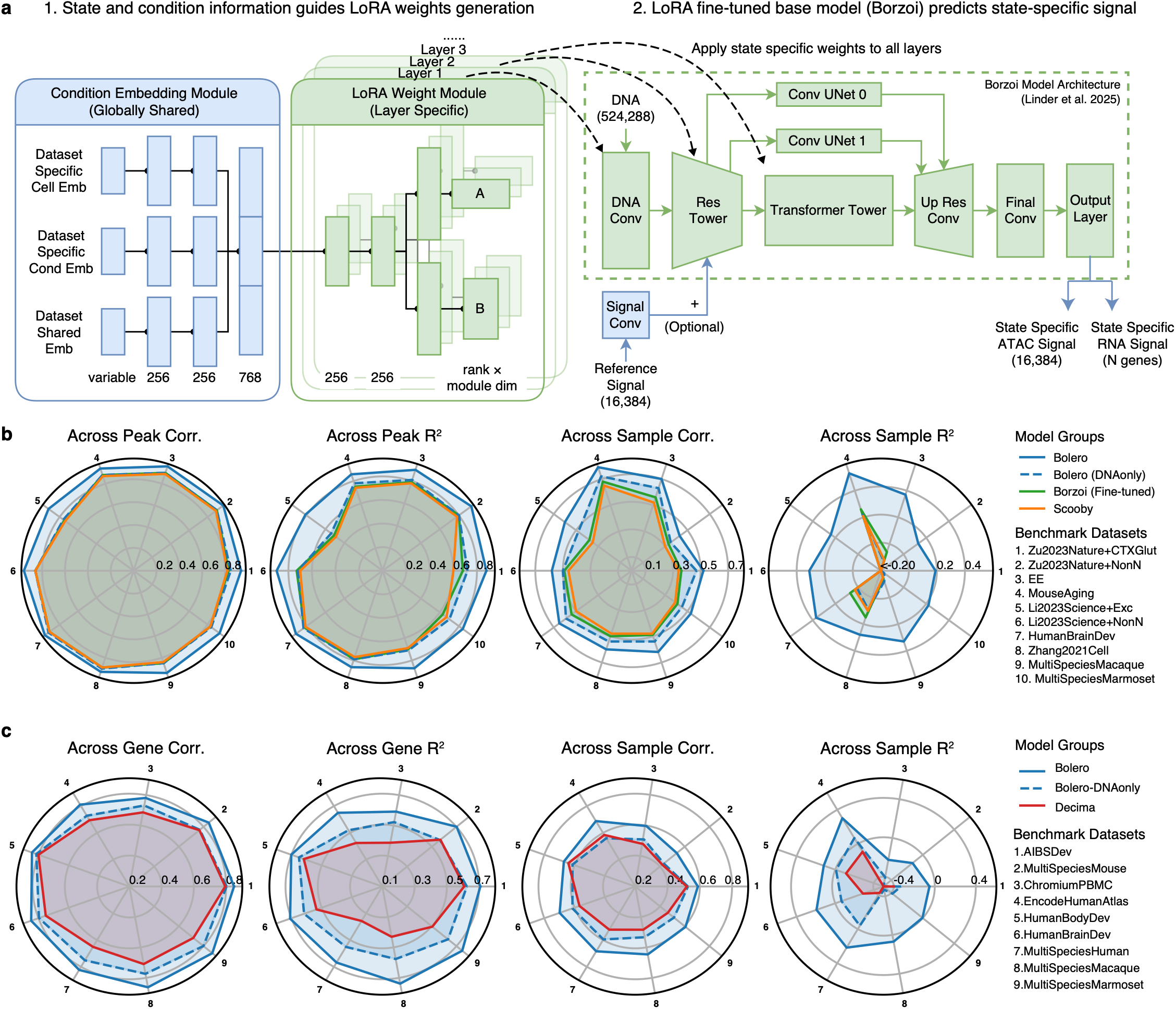
Model architecture and prediction-accuracy benchmarks. **a,** Architecture schematic: a globally-shared condition-embedding module turns dataset-specific cell and condition embeddings, together with a dataset-shared AP-1 activity score, into inputs for per-layer conditional low-rank (LoRA) weight modules, which generate state-specific weights applied across all layers of the Borzoi backbone (DNA, 524,288 bp; optional reference ATAC signal) to predict state-specific accessibility and gene expression. **b,c,** Radar plots of accessibility (b) or gene-expression (c) prediction accuracy across benchmark datasets for different models, over across-peak/gene and across-sample Pearson correlation and R². Bolero (DNA only) represents Bolero without bulk signal as input.

**Extended Data Figure 2.**
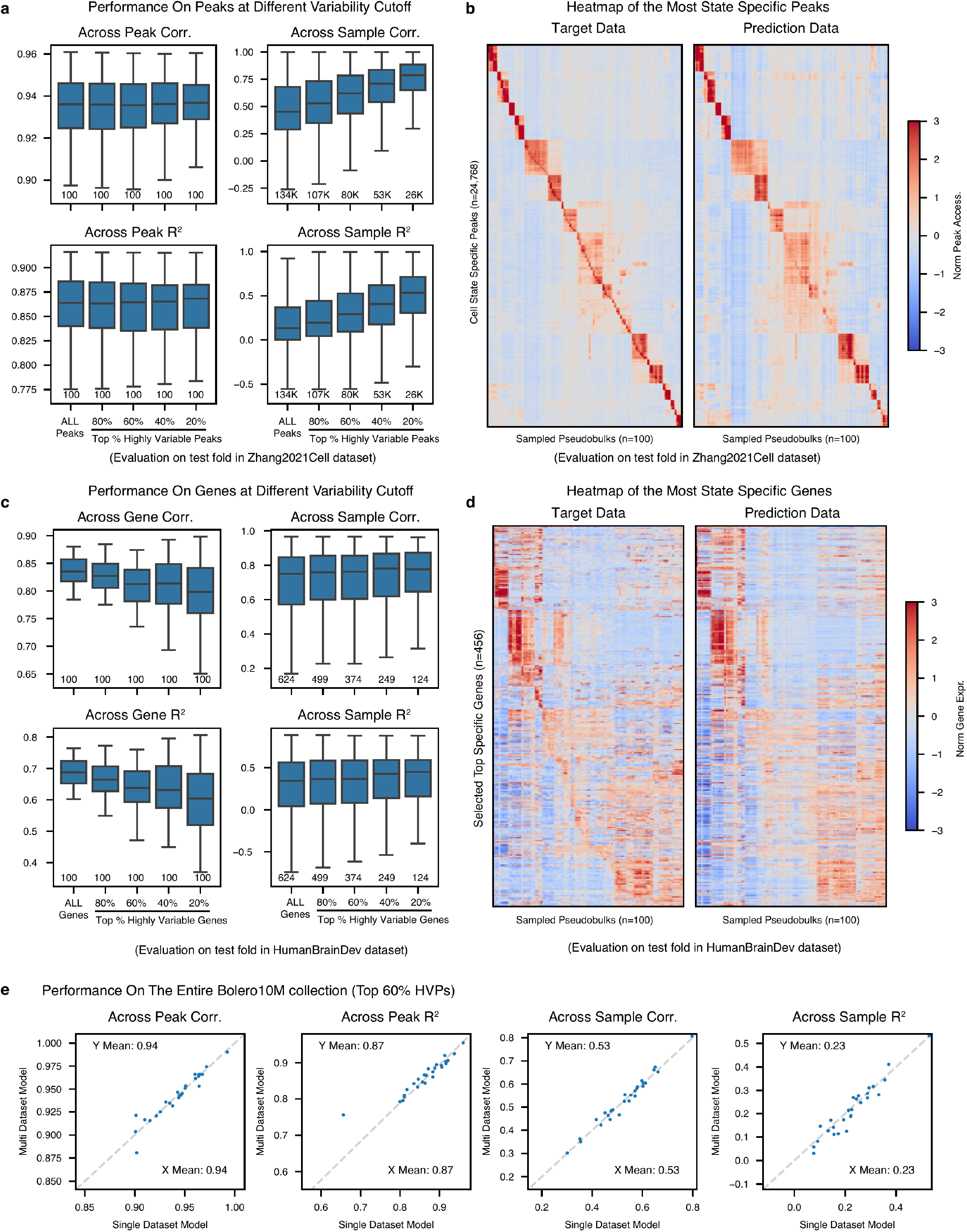
Prediction accuracy across variability cutoffs and state-specific reconstruction. **a,b,** Accessibility (a) or gene expression (b) prediction accuracy on Zhang2021Cell (a) or HumanBrainDev (b) across peak/gene variability cutoffs (ranked by standard deviations across pseudobulks), over across-peak/gene and across-sample Pearson correlation and R². Sample size is annotated under each box. Centre line denotes the median; box limits denote first and third quartiles; and whiskers denote 1.5 × the interquartile range. **c,d,** Measured (left) and predicted (right) accessibility (c) or gene expression (d) heatmaps of state-specific peaks (c) or genes (d). Among the top 40% most variable peaks (all genes), the 500 (50) with the highest fold-change over the mean of other states are taken per state and pooled. Columns (pseudobulks) are hierarchically clustered, and rows are grouped by the state each feature is specific to. Values are per-sample normalised counts after log1p and per-peak/gene z-score. **e,** Single-dataset versus multi-dataset Bolero accuracy (WholeHuman or WholeMouse, trained on all human or all mouse datasets) on Bolero-10M datasets (n = 27 dataset subsets from 18 human and mouse datasets) at top 60% variable peaks, over across-peak and across-sample Pearson correlation and R².

**Extended Data Figure 3.**
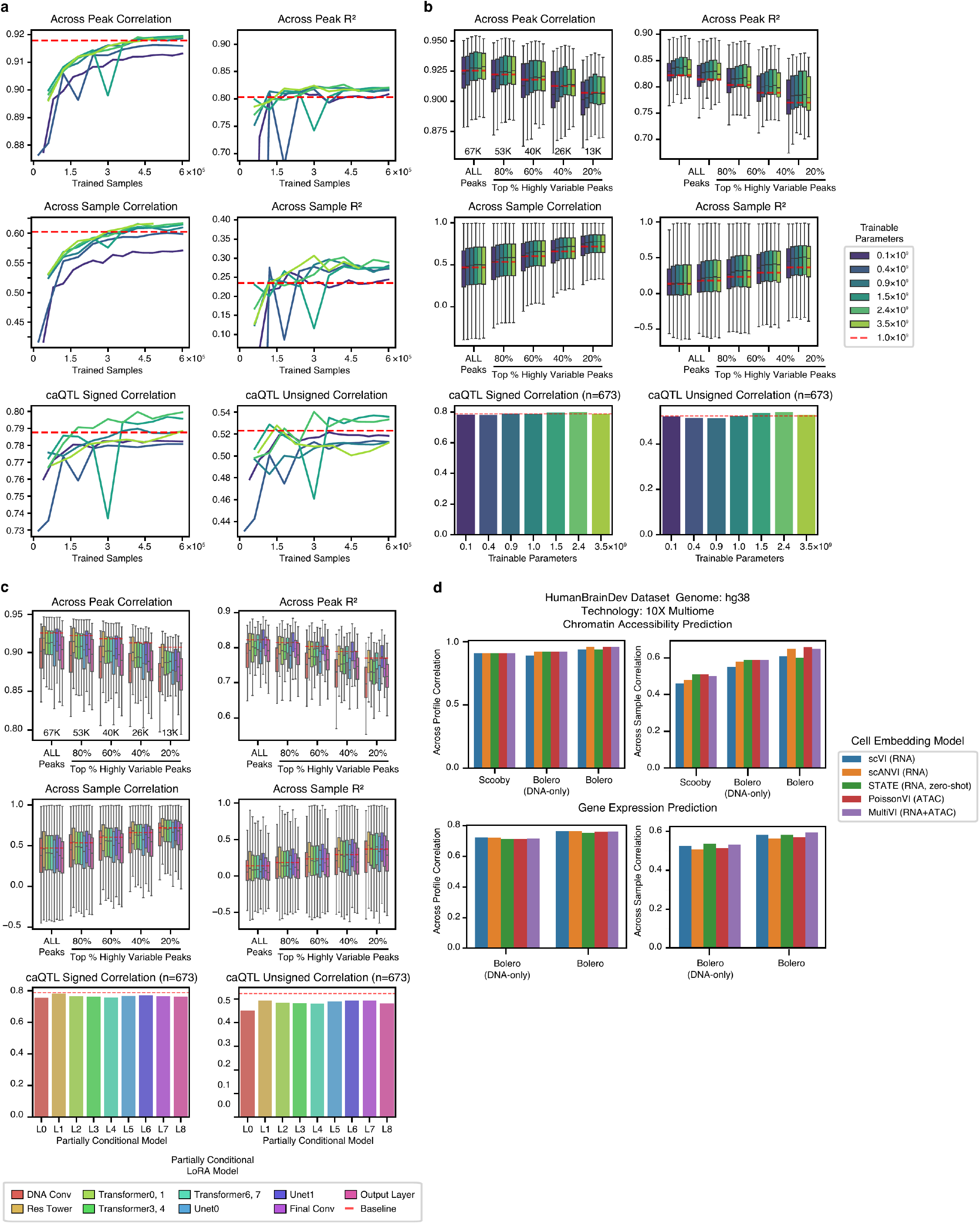
Model scaling, conditioning-layer ablation and embedding selection. **a,b,** Accuracy versus the number of trained samples (pseudobulk x genome region, x-axis in a), peak variability cutoffs (ranked by standard deviations across pseudobulks, x-axis in b), or the number of trainable parameters (colours) over across-peak (top) and across-sample (middle) correlation and R², and signed and unsigned caQTL correlation (bottom, n = 673). **c,** Accuracy of the full conditional model versus partially conditional models with conditioning in a single backbone module. **d,** Embedding benchmark on HumanBrainDev for ATAC (top) or RNA (bottom) predictions across cell-embedding methods for different models, over across-peak/gene and across-sample correlation. For all box plots, the centre line denotes the median; box limits denote first and third quartiles; and whiskers denote 1.5 × the interquartile range.

**Extended Data Figure 4.**
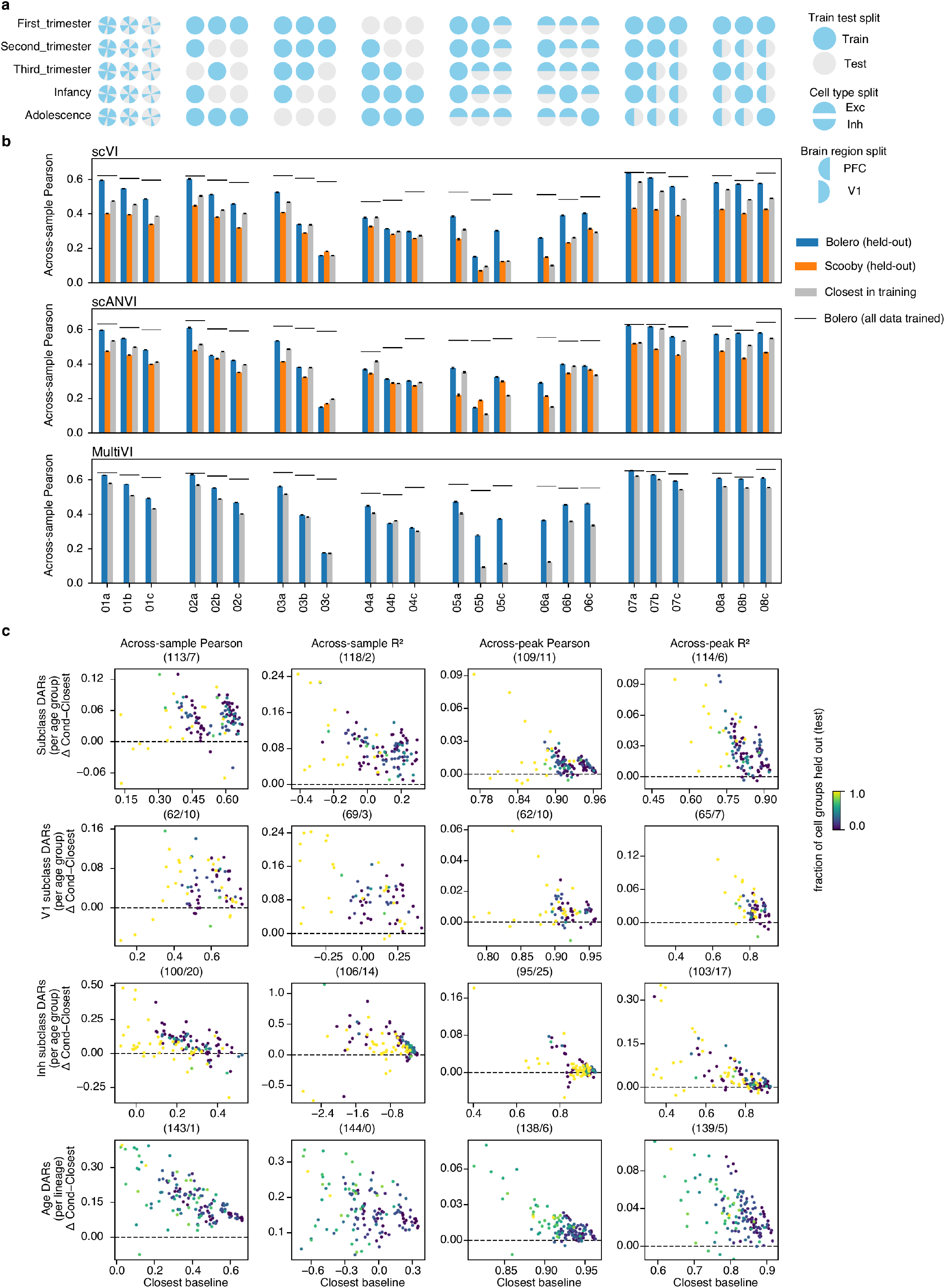
Held-out cell-state generalisation across embeddings and held-out axes. **a,** Train/test split per experiment. **b,** Across-sample Pearson across different models per case, for the scVI (top), scANVI (middle) and MultiVI (bottom) cell-state embeddings; Bolero beat the Closest baseline in 22 of 24 cases with scVI, 21 of 24 with scANVI and 23 of 24 with MultiVI. The black line is Bolero trained on all pseudobulks, and the bars are trained on training-only pseudobulks. The blue and orange bars are predictions using the condition and cell embedding of the test pseudobulk, with the orange (Scooby) bar in the scVI and scANVI rows only, and the grey bar is Bolero prediction using the closest training pseudobulk to the test. Error bars denote s.e.m. across peaks. **c,** Performance differences between Bolero and the closest (y-axis) versus the closest (x-axis) on different DARs (rows) over four metrics (columns). Each dot is a group of pseudobulks (age groups in rows 1-3, lineage groups in row 4) in each case (e.g. Case 01a - 3rd Trimester), coloured by the fraction of pseudobulks from the group held out in the case. Numbers in subplot titles represent the number of dots above/below the horizontal zero line (Bolero performs the same as closest).

**Extended Data Figure 5.**
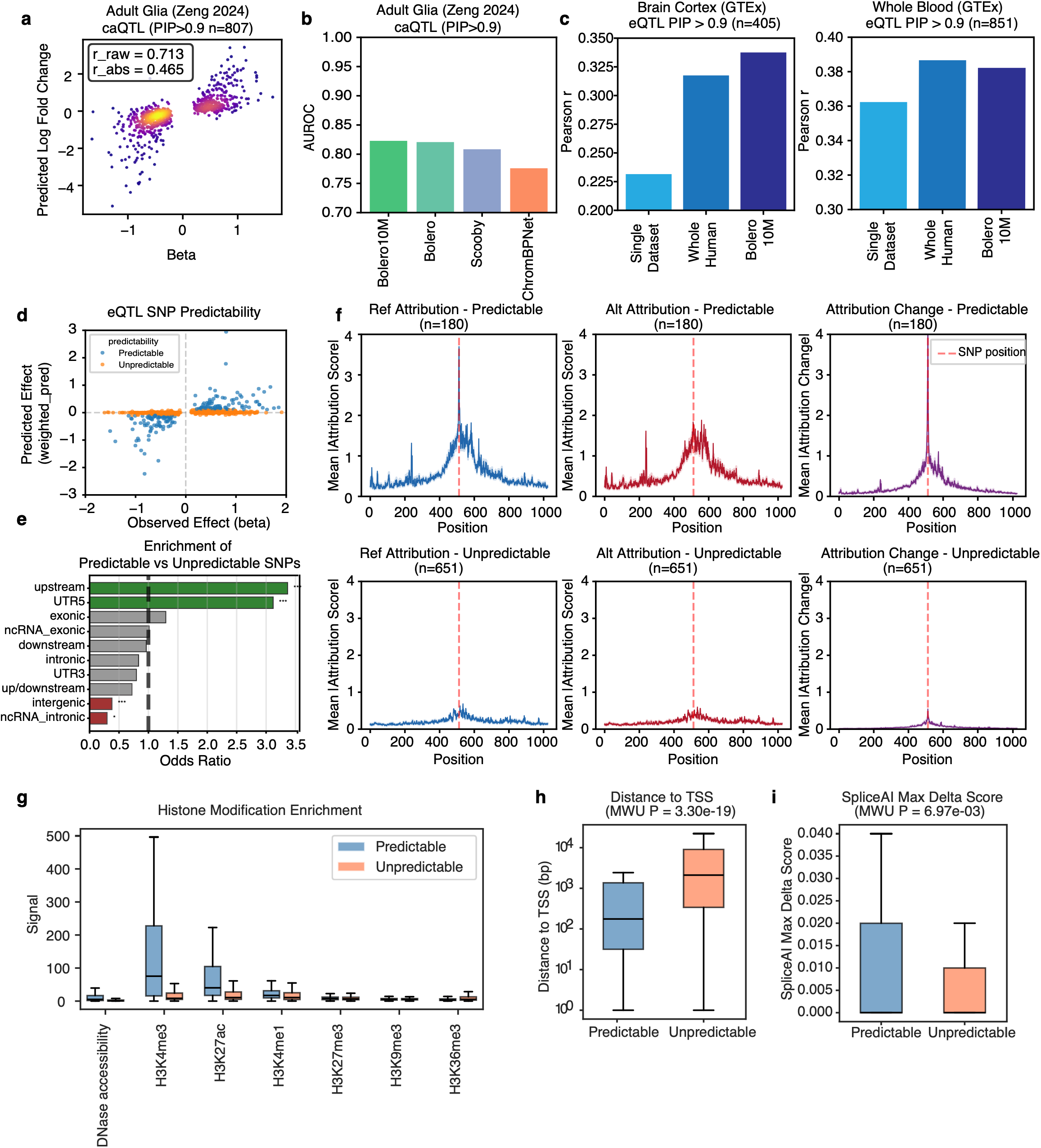
Determinants of variant-effect predictability and model-scale comparison. **a,** Observed versus predicted adult-glia caQTL effects (Zeng 2024, PIP > 0.5, n = 807; Pearson r = 0.71 on signed effects and r = 0.47 on absolute effect sizes). **b,** AUROC for prioritising fine-mapped adult-glia caQTLs, for Bolero-10M, Bolero (brain only), Scooby and ChromBPNet. **c,** Pearson correlation between real and predicted eQTL effect sizes across models trained on different dataset scales for brain-cortex (GTEx, PIP > 0.9, n = 405) and whole-blood (n = 851). Single dataset represents MultiSpeciesHuman for cortex and ChromiumPBMC for blood; whole human is trained on all human datasets and Bolero-10M on the full atlas. **d,** Observed versus predicted effect of predictable (correctly signed prediction; n = 180) and unpredictable (n = 651) variants. **e,** Odds-ratio enrichment of predictable (n = 180) over unpredictable (n = 651) eQTLs across genomic categories. **f,** Mean absolute per-base reference (left), alternate (middle), and change (right) attribution profiles across the 1-kb window centred on the predictable (top, n = 180) and unpredictable (bottom, n = 651) eQTLs; the 17 PBMC pseudobulk tracks of each eQTL were averaged first; shading, s.e.m. across eQTLs. **g–i,** Active/repressive histone-mark and accessibility signal (g), distance to the nearest TSS (h), and SpliceAI maximum delta score (i) of predictable (n = 178 (g), 178 (h), 103 (i)) and unpredictable (n = 635 (g), 635 (h), 364 (i)) eQTLs. Only eQTLs with annotations are shown in g and h, and eQTLs with SpliceAI predictions are shown in i. For all box plots, the centre line denotes the median; box limits denote first and third quartiles; and whiskers denote 1.5 × the interquartile range.

**Extended Data Figure 6.**
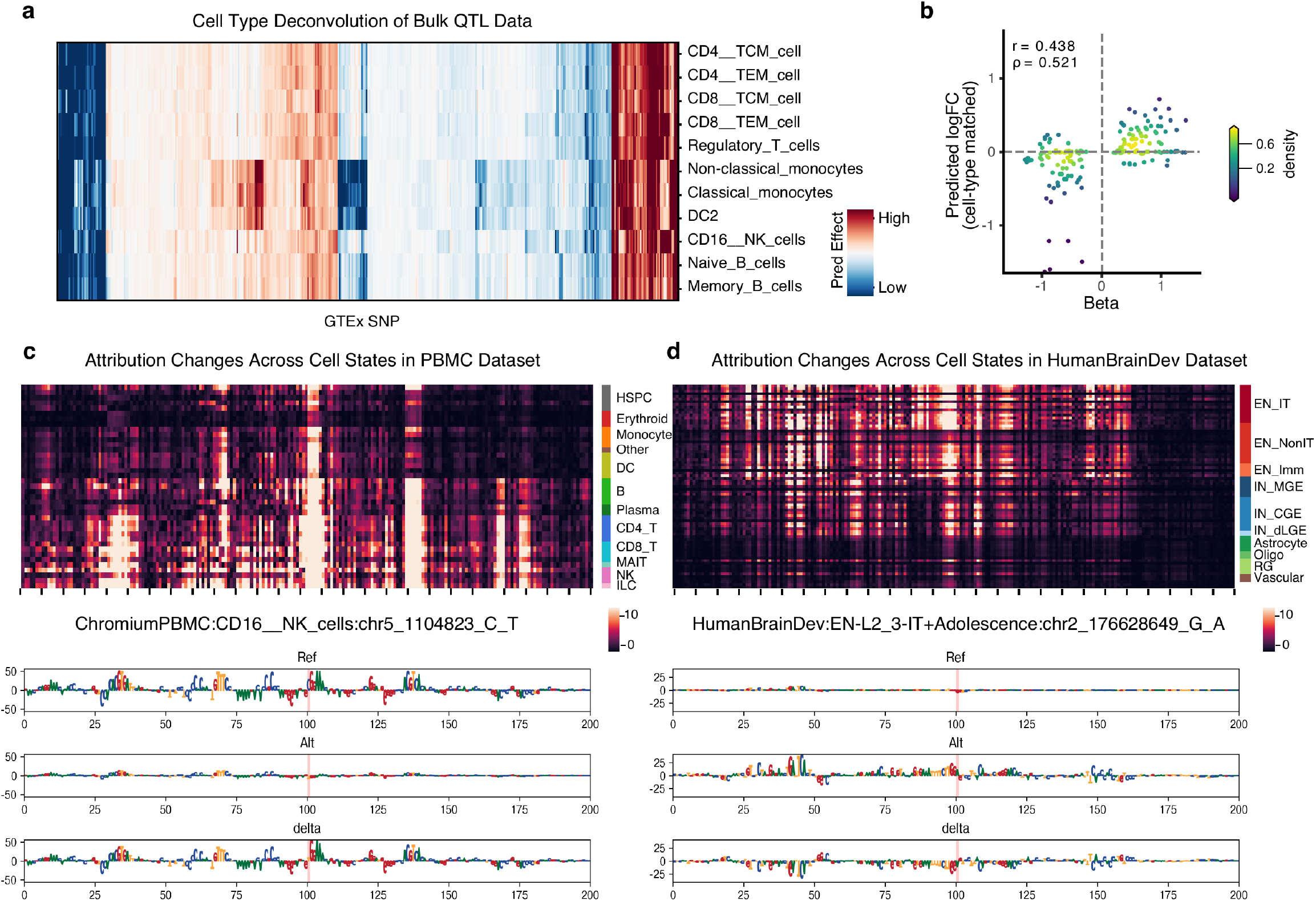
Bulk-QTL deconvolution and atlas-wide single-variant attribution. **a,** Heatmap of Bolero’s predicted whole-blood GTEx eQTL effect deconvolved into 11 PBMC cell types (rows). Only eQTLs with a non-negligible predicted effect are shown (n= 437, columns, hierarchically clustered). **b,** For whole-blood GTEx eQTLs overlapping OneK1K cell-type-specific eQTLs, measured cell-type-specific effect size (beta) in OneK1K versus Bolero’s cell-type-matched predicted log fold-change. Colour is point density, and each point is a cell-type × eQTL-SNP pair (n = 178 cell-type × eQTL pairs from 73 gene-variant pairs fine-mapped in OneK1K, PIP > 0.9, overlapping the GTEx set; Pearson r = 0.44, Spearman ρ = 0.52). **c,d,** Single-variant attribution change for a blood (c) or cortex (d) eQTL across a 200-bp window centred on the variant. Top: Heatmap of the predicted per-base attribution change across pseudobulks (rows) grouped by cell class. Bottom: DNA sequence across the same window with letter height representing reference-allele, alternate-allele and delta (reference − alternate) per-base attribution, with the variant position highlighted.

**Extended Data Figure 7.**
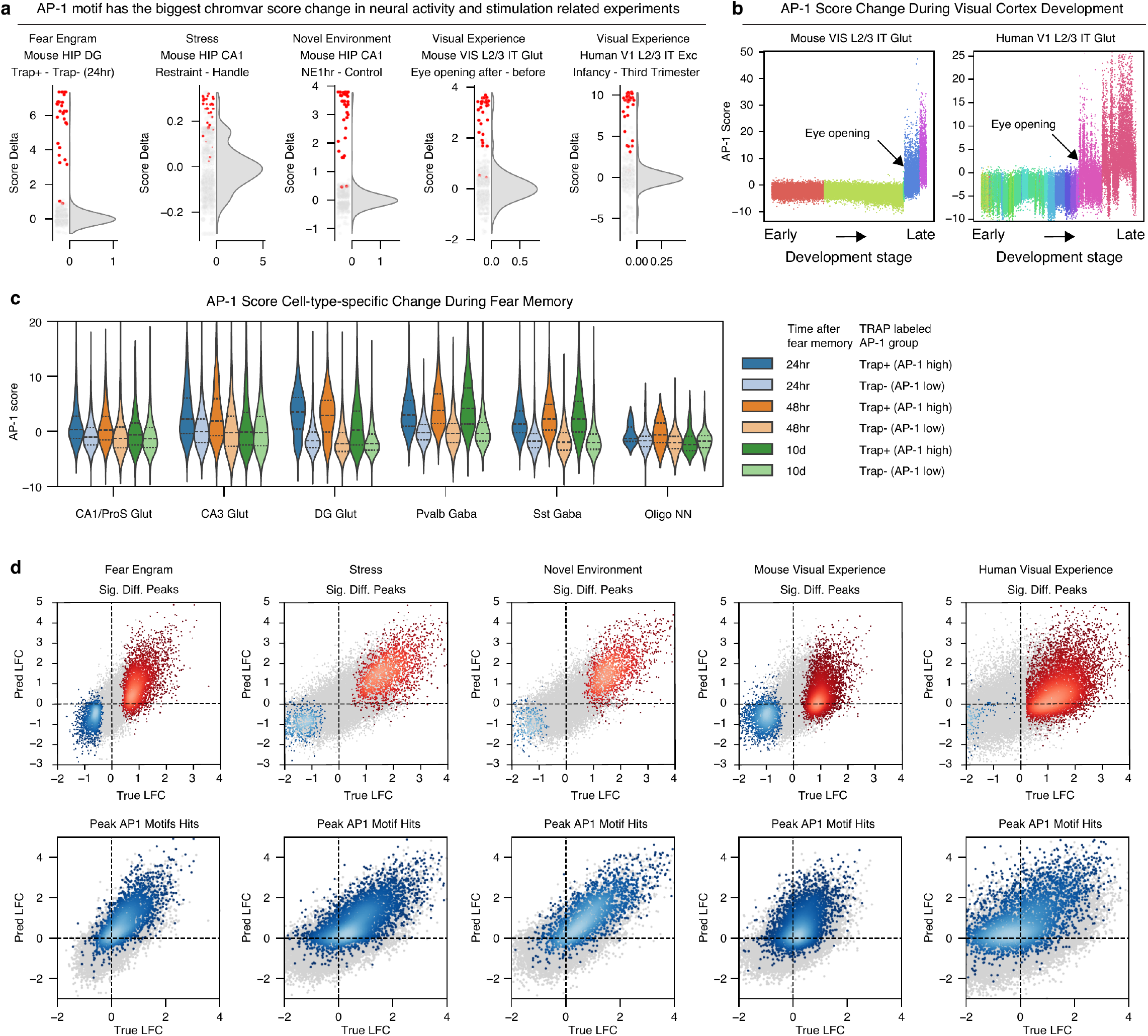
AP-1 activity score across stimulation paradigms. **a,** Per-motif chromVAR score changes across five paradigms as a half-violin density plus jittered strip of motifs. AP-1 motifs are highlighted in red. **b,** Per-cell AP-1 chromVAR score along the visual-cortex developmental trajectory, coloured by developmental age, for mouse VIS and human V1 L2/3 IT. **c,** Per-cell AP-1 chromVAR score violins by subclass and experimental group across hippocampal cell types. **d,** Observed versus predicted log fold-change at peaks for each of the five paradigms coloured by density. Top: significant differential peaks in red (up) or blue (down), non-significant peaks grey. Bottom: peaks with an AP-1 motif in blue, without in grey.

**Extended Data Figure 8.**
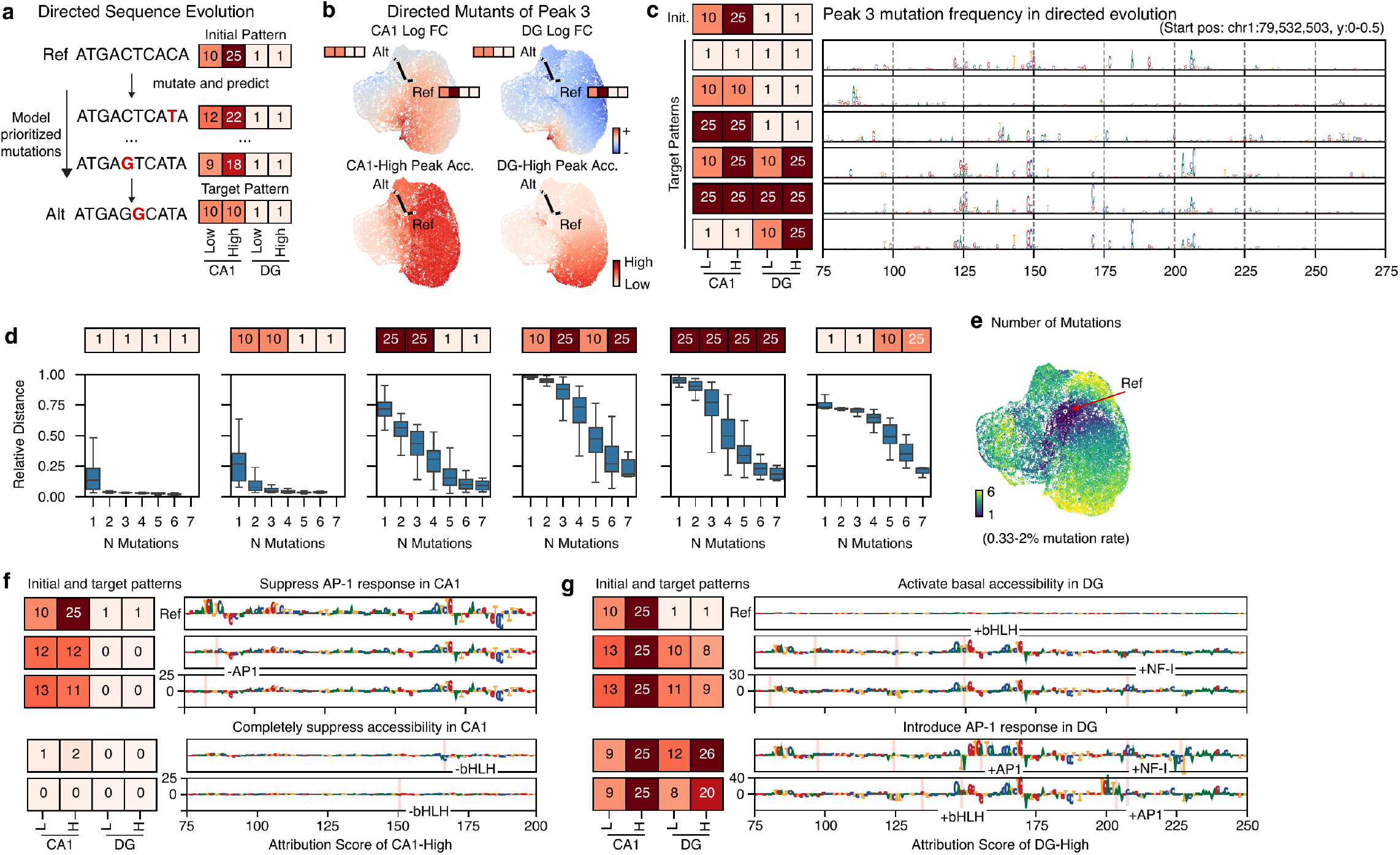
Enhancer design by directed evolution. **a,** Schematic of directed sequence evolution. The state of a sequence is represented as a four-dimensional vector of predicted accessibility in CA1 and DG at AP-1-low and AP-1-high. **b,** UMAP of all generated directed-mutant sequences (n = 29,737) from an example peak (Peak 3 in Fig. 4e) to eight target states, coloured by predicted CA1 log fold-change, DG log fold-change, CA1-high accessibility and DG-high accessibility. Reference (Ref) and an example evolved outcome (Alt) sequence with the evolution path are labelled. **c,** DNA sequence of the example peak with letter height representing mutation frequency to achieve each directed-evolution target pattern. **d,** Relative distance to the design target (normalised to the reference) versus the number of mutations per design (n = 48,000, 50 experiments x 15 rounds x 64 candidate sequences), for each of six target patterns. Centre line denotes the median; box limits denote first and third quartiles; and whiskers denote 1.5 × the interquartile range. **e,** The same directed-mutant embedding as in b, coloured by the number of mutations per design. **f,g,** DNA sequences of different evolved Peak 3 that lose or gain peak accessibility or AP-1 response in specific cell type, with letter height representing the attribution score.

**Extended Data Figure 9.**
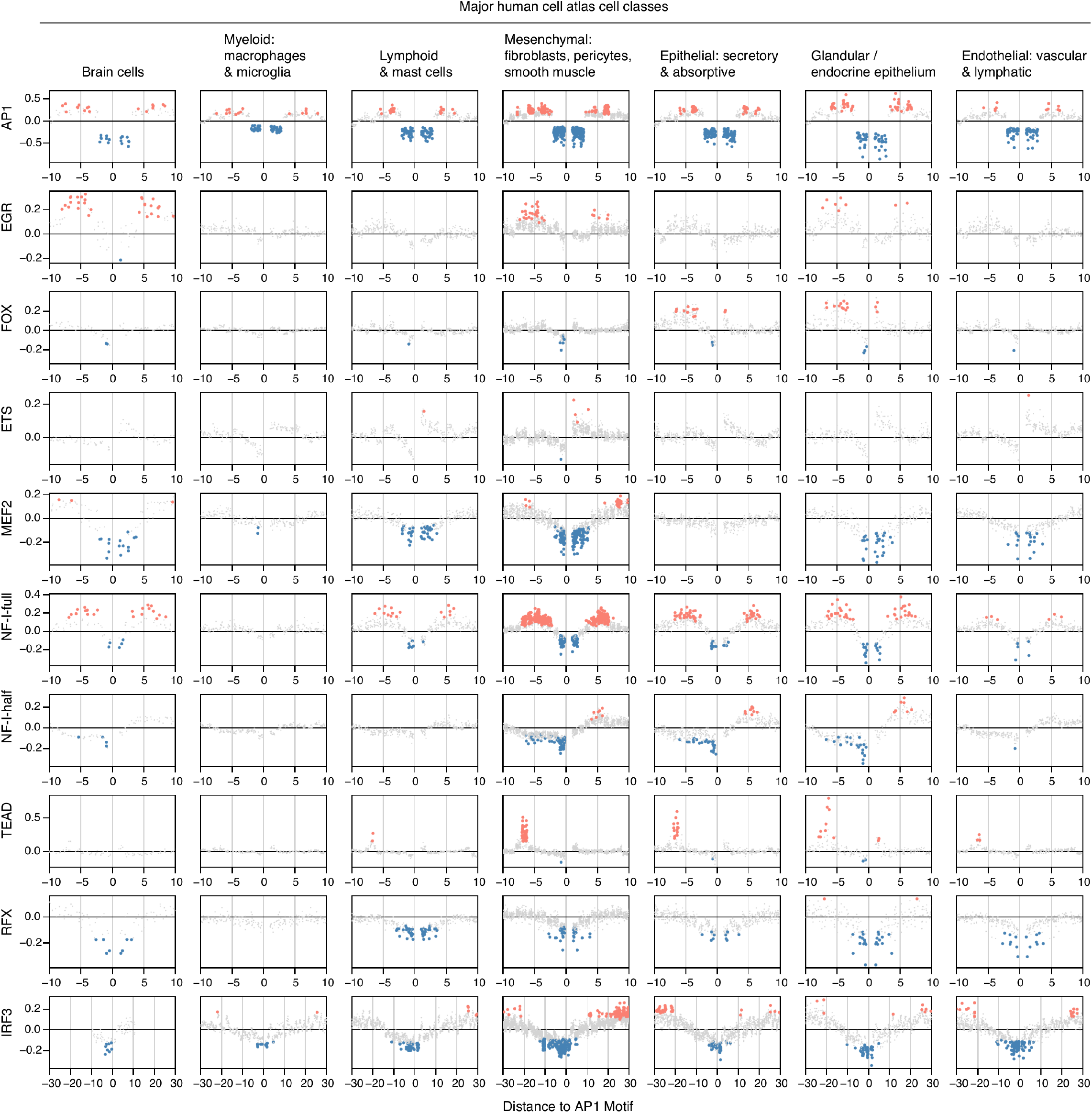
Motif-insertion assay across human-atlas cell classes. Predicted AP-1 effects (log2 fold-change) versus the signed distance between an inserted AP-1 and an inserted partner motif (rows), shown separately for the seven major cell classes of the human whole-body atlas (Zhang 2021; columns). Each dot is a pseudobulk × spacing, coloured by the sign of a significant effect and grey otherwise, with significant dots drawn larger.

**Extended Data Figure 10.**
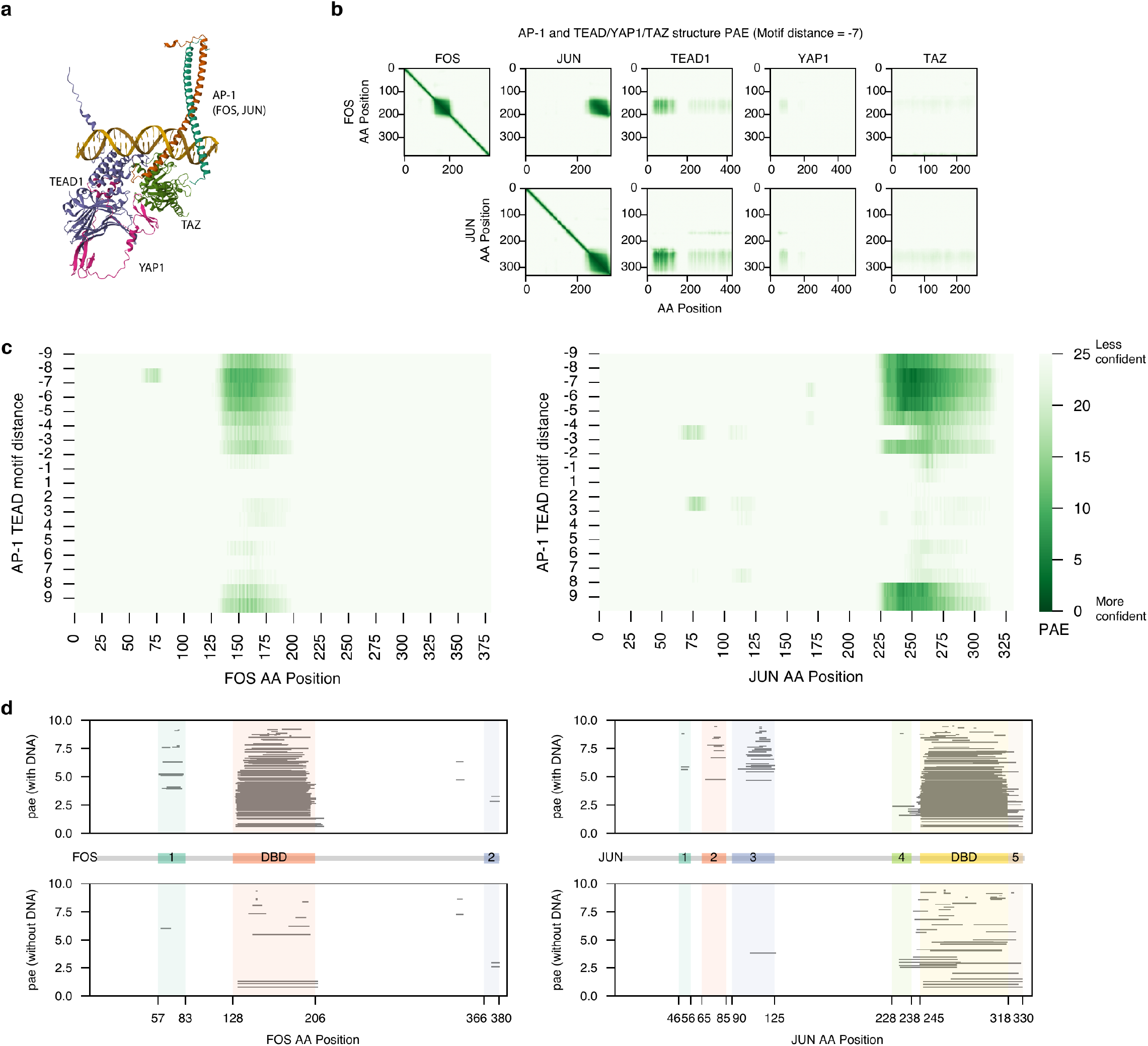
AlphaFold3 structural analysis of AP-1–cooperator co-binding. **a,** AlphaFold3 model of AP-1 (FOS, JUN) with TEAD1, YAP1 and TAZ on DNA (chains coloured; DNA in gold). **b,** Grid of inter-chain predicted-aligned-error (PAE) heatmaps of FOS (top) or JUN (bottom) amino-acid position against each chain (FOS, JUN, TEAD1, YAP1, TAZ), for the complex modelled with the AP-1 and TEAD motifs separated by −7 bp (darker = more confident). **c,** Inter-chain PAE between FOS (left) or JUN (right) and TEAD across AP-1–TEAD motif distance and FOS/JUN amino-acid position. **d,** Confident inter-chain interaction segments for FOS (left) and JUN (right) with DNA (top) and without DNA (bottom), plotted by amino-acid position; the middle track maps FOS/JUN protein domains including the bZIP DNA-binding domain (DBD).

**Extended Data Figure 11.**
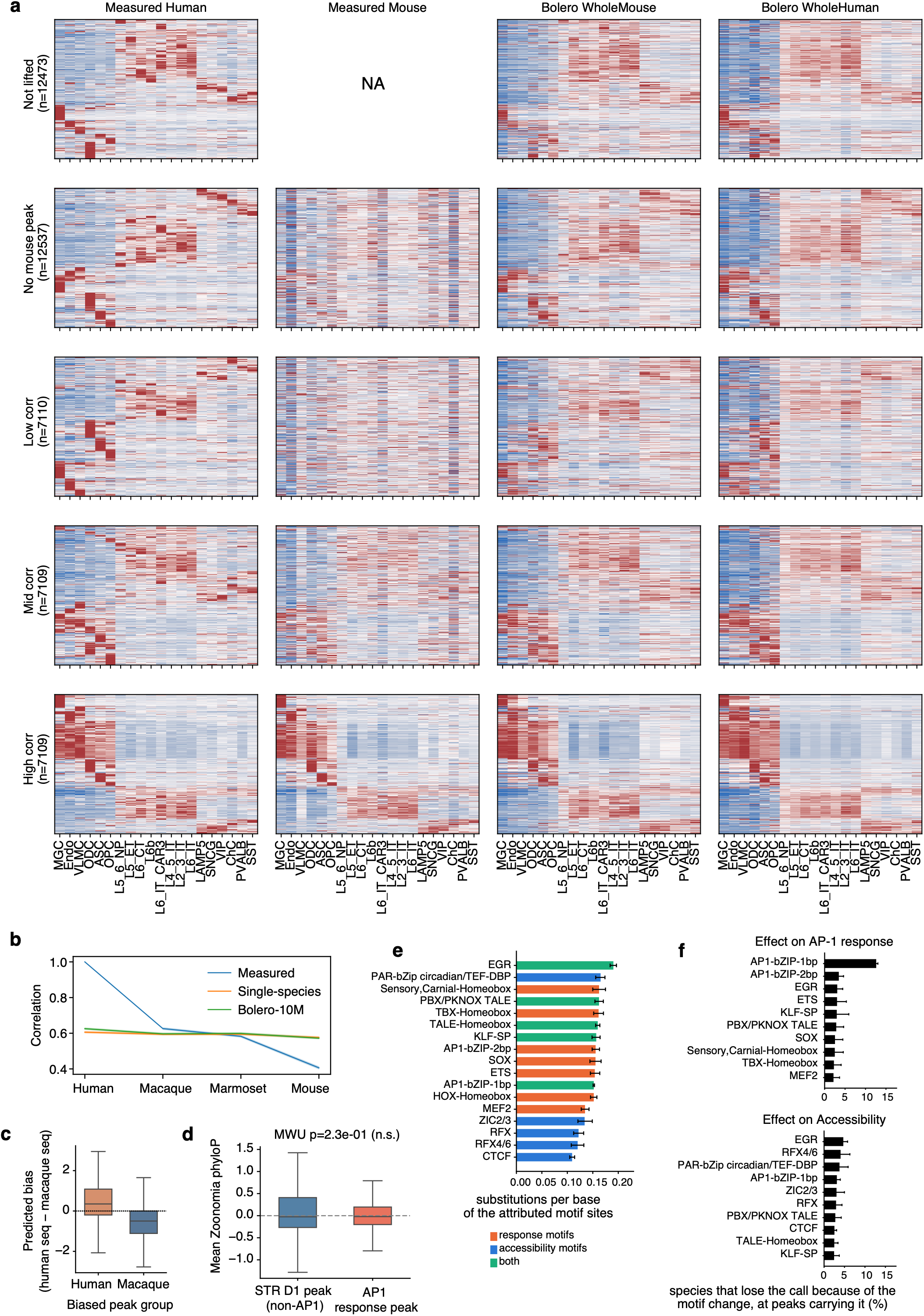
Cross-species conservation of predicted accessibility. **a,** Heatmap of measured accessibility in human (Measured Human), in mouse at liftover coordinates (Measured Mouse; NA for not-lifted peaks), or predicted with human sequence by Bolero model trained across mouse (Bolero WholeMouse) or human (Bolero WholeHuman) datasets, stratified into five conservation-group blocks of subclass DARs (row, Fig. 6d). Values are z-scored per peak across subclasses. **b,** Correlation across the 20 cortical subclasses between measured human accessibility and measured data from other species, or predicted data by the single-species models and Bolero-10M. **c,** Predicted bias (human sequence minus macaque sequence) for the human-biased and macaque-biased peak groups. **d,** Zoonomia phyloP (241-way mammalian) for AP-1-responsive (n = 15,120) vs other STR_D1 accessible peaks (n = 208,718). **e,** Cross-species substitution rates at attributed motif sites, expressed as substitutions per base for the indicated motifs. Motifs were the union of those in Fig. 6m and panel f. **f,** Percentage of species predicted to lose predicted AP-1 response at AP-1-responsive D1 peaks (n = 8,769, top) or chromatin accessibility at non-AP-1-responsive D1 peaks (n = 5,321, bottom) owing to changes at attributed motif sites. Values are computed only on peaks carrying the motif. The ten motifs with the largest contributions are shown. Error bars are 95 percent bootstrap intervals over peaks, from 1,000 draws.

